# Combinatorial regulation of the balance between dynein microtubule end accumulation and initiation of directed motility

**DOI:** 10.1101/126508

**Authors:** Rupam Jha, Johanna Roostalu, Martina Trokter, Thomas Surrey

**Affiliations:** The Francis Crick Institute, 1 Midland Road, London NW1 1AT, United Kingdom

## Abstract

Cytoplasmic dynein is involved in a multitude of essential cellular functions. Dynein’s activity is controlled by the combinatorial action of several regulators. The molecular mechanism of this regulation is poorly understood. Using purified proteins, we reconstitute the regulation of the human dynein complex by three prominent regulators on dynamic microtubules in the presence of end binding proteins (EBs). We find that dynein can be in biochemically and functionally distinct pools: either passively tracking dynamic microtubule plus-ends in an EB-dependent manner or moving processively towards minus ends in an adaptor protein-dependent manner. Whereas both dynein pools share the dynactin complex, they have opposite preferences for binding other regulators, either the adaptor protein Bicaudal D2 (BicD2) or the multifunctional regulator Lisencephaly-1 (Lis1). Remarkably, dynactin, but not EBs, strongly biases motility initiation locally from microtubule plus ends by autonomous plus end recognition. BicD2 and Lis1 together control the overall efficiency of motility initiation. Our study provides insight into the mechanism of dynein activity regulation by dissecting the distinct functional contributions of the individual members of a dynein regulatory network.

## INTRODUCTION

Cytoplasmic dynein-1 (here called dynein) is the major minus-end directed microtubule motor in most eukaryotes (Vale, 2003). It is involved in a variety of cellular functions ranging from organelle transport (Allan, 2011) to spindle pole focussing (Merdes et al., 2000), removal of checkpoint proteins from kinetochores (Griffis et al., 2007), and spindle positioning (McGrail and Hays, 1997). Dynein is a 1.4 MDa protein complex consisting of two copies of six different subunits that assembled into a tail domain from which two motor domains protrude (Pfister et al., 2006; Vale, 2003; Vallee et al., 1988). The motile properties of metazoan dynein depend strongly on its interaction with a variety of regulatory proteins whose mechanisms of action and their combinatorial interplay are poorly understood (Cianfrocco et al., 2015; Vallee et al., 2012).

The major regulator of dynein that is required for most of its cellular functions, is dynactin, a 1.5 MDa protein complex that consists of different copies of twelve different protein subunits (Karki and Holzbaur, 1999; Schroer, 2004; Urnavicius et al., 2015). Despite being the key dynein regulator, dynactin itself interacts only weakly with dynein (McKenney et al., 2014; Schlager et al., 2014). Additional adaptor proteins that recruit dynein to membranes of cargoes, stabilise this interaction, leading to ternary complex formation and activation of processive dynein motility (McKenney et al., 2014; Schlager et al., 2014). Ternary complex formation is thought to release dynein from its autoinhibited state, possibly by separating the two motor domains (Carter et al., 2016; Chowdhury et al., 2015; Urnavicius et al., 2015).

There are several such adapter proteins that link dynein/dynactin to different cargos like organelles and kinetochores or to the cell cortex (Kardon and Vale, 2009; McKenney et al., 2014). One example is the metazoan-specific adapter protein Bicaudal-D2 (BicD2) that is critical for bidirectional transport of mRNA particles, and contributes to the positioning of the endoplasmatic reticulum, the Golgi apparatus and the nucleus (Bullock and Ish-Horowicz, 2001; Hoogenraad and Akhmanova, 2016). The N-terminal coiled-coil of BicD2 mediates the interaction between the dynein tail and dynactin, which is crucial for processive minus end directed dynein motion (Chowdhury et al., 2015; McKenney et al., 2014; Schlager et al., 2014; Urnavicius et al., 2015).

Despite being a minus-end directed motor, dynein is also well known to accumulate at the plus ends of growing microtubules in cells (Kardon and Vale, 2009). This interaction is thought to enable dynein to initiate cargo transport at microtubule plus ends (Egan et al., 2012; Moughamian and Holzbaur, 2012; Vaughan et al., 2002) and to facilitate dynein-mediated interactions between microtubule plus ends and other cellular structures like the cell cortex (McGrail and Hays, 1997) or the kinetochores (Faulkner et al., 2000). Different pathways are known to be responsible for microtubule plus end accumulation of dynein. Kinesin-dependent transport has been observed in fungi and neurons of metazoans, whereas EB1 family protein (EB)-dependent end tracking constitutes the dominant pathway in non-neuronal cultured mammalian cells (Carvalho et al., 2004; Duellberg et al., 2014; Moughamian et al., 2013; Roberts et al., 2014).

EBs autonomously bind growing microtubule plus end regions by recognising the nucleotide state of freshly added tubulins (Akhmanova and Steinmetz, 2015; Bieling et al., 2007; Maurer et al., 2011). EBs recruit a multitude of other plus-end tracking proteins (Akhmanova and Steinmetz, 2015), including dynactin which is additionally required for plus-end tracking of dynein in cells (Dixit et al., 2008; Ligon et al., 2003; Vaughan et al., 1999). The critical dynactin subunit for its plus end-tracking behaviour is p150^Glued^ (called p150 here). Its N-terminal CAP-Gly domain protrudes from the shoulder of the dynactin complex (Chowdhury et al., 2015; Urnavicius et al., 2015) and binds directly to microtubules (Culver-Hanlon et al., 2006; Ross et al., 2006), as well as to EBs (Honnappa et al., 2006). In vitro reconstitution experiments with a p150 fragment showed that the p150 CAP-Gly domain and the first coiled-coil of p150 which interacts with the intermediate chain of the dynein complex (Karki and Holzbaur, 1995; King et al., 2003; Vaughan and Vallee, 1995), were sufficient for mediating EB-dependent end tracking of the human dynein complex (Duellberg et al., 2014). The regulation of the balance between dynein microtubule end tracking and its processive motility could however not be studied in these experiments as this requires the presence of the entire dynactin complex. Furthermore, in a recent cryo-electron microscopy structure of the dynactin complex, both the p150 CAP-Gly domain and the first coiled-coil appear buried in the groove of the dynactin shoulder, raising the question of whether p150 in the context of the entire dynactin complex can mediate EB-dependent dynein end tracking at all, or if additional factors might be needed to release a potential autoinhibition of dynactin (Urnavicius et al., 2015).

Dynactin-dependent plus-end localisation of dynein has been demonstrated to be involved in the control of transport initiation in distal neurites (Moughamian et al., 2013; Nirschl et al., 2016). In non-polarised cells, the role of dynactin in transport initiation is however not so well understood (Dixit et al., 2008; Kim et al., 2007). From a mechanistic point of view, it is unclear whether EB-recruited dynactin can contribute to initiating transport, thereby promoting transport initiation preferentially from microtubule plus ends. Alternatively, it is conceivable that dynein tracking microtubule ends and dynein initiating processive runs belong to separate dynein pools.

Another major question concerns the mechanism by which the dynein regulator Lissencephaly 1 (Lis1) controls the balance between plus-end tracking of dynein versus initiation of minus-end directed motility. Lis1 has been reported to regulate both initiation minus-end directed motility (Egan et al., 2012; Moughamian et al., 2013; Splinter et al., 2012) and plus-end tracking of dynein (Coquelle et al., 2002; Splinter et al., 2012) in cells.

However, Lis1 was not required for dynein end tracking in a minimal in vitro reconstitution (Duellberg et al., 2014), raising the question as to why Lis1 is crucial for dynein end tracking in cells. Lis1 is a homodimeric 45 kDa protein that binds directly to the dynein motor domain (Kardon and Vale, 2009; Mateja et al., 2006), reported to induce a more strongly microtubule-bound state of metazoan dynein (McKenney et al., 2010; Torisawa et al., 2011; Yamada et al., 2008), thereby increasing the force produced by dynein (McKenney et al., 2010; Reddy et al., 2016), and slowing down microtubule transport by surface immobilised dynein motors (Torisawa et al., 2011; Wang et al., 2013; Yamada et al., 2008).

Our understanding of how the combined action of various dynein regulators controls dynein behaviour is limited. This is at least in part due to the lack of in vitro reconstitutions allowing the simultaneous study of processive motility and microtubule plus end tracking of dynein that would allow the dissection of the respective roles of distinct regulators.

Here, we perform in vitro reconstitutions probing the mechanism of combinatorial regulation of recombinant human dynein by three human dynein regulators - dynactin, BicD2 and Lis1 - in the presence of EB proteins. We find that the dynactin complex alone can mediate EB-dependent end tracking of dynein, thereby biasing dynein localisation towards microtubule ends. Despite competition for dynein binding, Lis1 and BicD2 cooperate to initiate processive motility. Plus end-tracking dynein and processively moving dynein seem to comprise two distinct dynein pools defined by distinct interaction partners. Finally, we find that dynactin has the previously unknown capacity to initiate processive dynein motility preferentially from microtubule plus ends, independent of EB proteins. These reconstitutions provide insight into the basic molecular mechanism of biased transport initiation of dynein from microtubule ends.

## RESULTS

To investigate the behaviour of human cytoplasmic dynein and dynactin on dynamic microtubules, we purified both protein complexes. GFP-tagged human dynein was expressed and purified from insect cells as described (Schlager et al., 2014; Trokter et al., 2012) (Methods). The human dynactin complex was purified from HeLa S3 cell cultures using a new method based on BicD2 affinity chromatography followed by ion exchange chromatography (Methods). Purity of both complexes was demonstrated by SDS PAGE (Fig. EV1A-C) and the subunit composition of the dynactin complex was verified by mass spectrometry (Fig. EV1A). To be able to study simultaneously dynamic microtubule end tracking and processive motility initiation of dynein/dynactin, we adjusted the conditions of previous in vitro reconstitution experiments, using here a buffer that was intermediate in ionic strength compared to previous dynein motility studies on static microtubules (McKenney et al., 2014; Schlager et al., 2014; Trokter et al., 2012) and microtubule end tracking experiments (Duellberg et al., 2014; Roberts et al., 2014) (Methods).

We first investigated whether the purified human dynactin complex can recruit dynein to dynamic microtubule plus ends in the presence of purified EB1 (Fig. EV1B). These experiments were intended to test if in the absence of other dynein/dynactin regulators the N-terminal part of the dynactin component p150 is undocked from the full dynactin complex and available for EB binding, and if the reported weak interaction between dynein and dynactin in the absence of cargo adaptors (Chowdhury et al., 2015; Urnavicius et al., 2015) can support efficient dynein microtubule end tracking. Using time-lapse dual colour total internal reflection fluorescence (TIRF) microscopy, we observed dynamic microtubules growing from immobilised seeds in the presence of 10 nM GFP-dynein, 20 nM dynactin, 20 nM purified EB1 and 17.5 μΜ Alexa568-tubulin. GFP-dynein accumulated strongly on growing microtubule plus and minus ends (Fig. 1A, B and Movie 1), typical for recruitment depending on EB1 family proteins (EBs). Additionally, some diffusive and static GFP-dynein binding events were observed all along the microtubules. GFP-dynein was also enriched on shrinking microtubule ends after microtubules had switched to depolymerisation (Fig. 1A).

**Figure 1.**
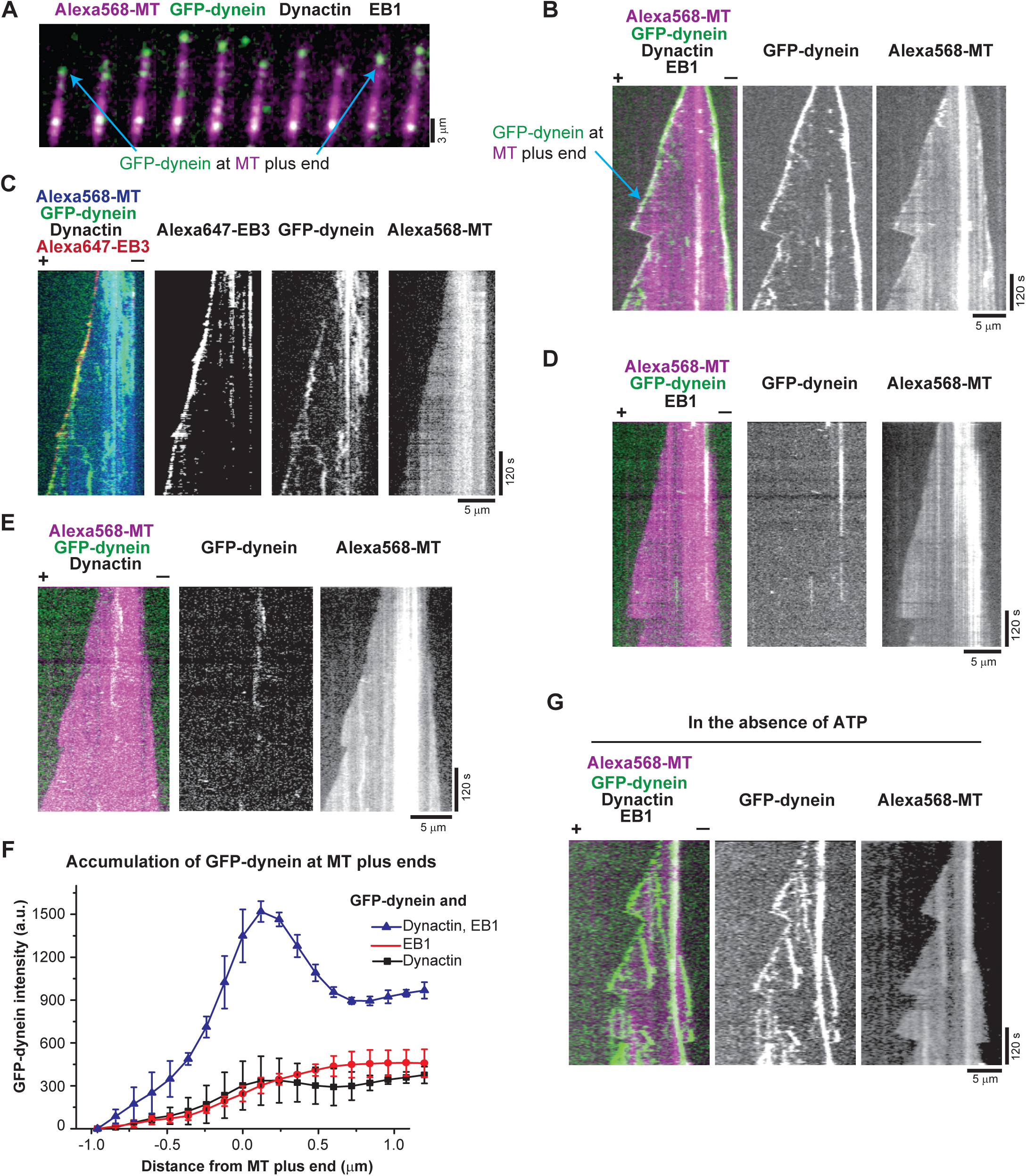
Microtubule plus-end tracking of dynein in the presence of dynactin and EB protein. (**A**) Time series of TIRF microscopy images and (**B**) individual and dual colour kymographs showing GFP-dynein (green in merge) localising to the plus ends of dynamic Alexa568-microtubules (Alexa568-MT, magenta in merge). Protein concentrations were 10 nM GFP-dynein, 20 nM human dynactin, 20 nM EB1 and 17.5 μΜ Alexa568-tubulin (5% labelling ratio). (**C**) Merged triple colour and single fluorescence channel kymographs showing microtubule end tracking of GFP-dynein (green in merge) and Alexa647-EB3 (red in merge) on dynamic Alexa568-microtubules (blue in merge). Concentrations were 10 nM GFP-dynein, 10 nM human dynactin, 10 nM Alexa647-EB3, and 17.5 μM tubulin. (**D**, **E**) Dual and single colour kymographs of GFP-dynein (green) on dynamic Alexa568-microtubules (magenta) in the absence of either dynactin (D) or EB proteins (E). Concentrations of the proteins present as in A. (**F**) Averaged fluorescence intensity profiles of GFP-dynein at growing microtubule plus ends in the presence of both human dynactin and EB1 (blue triangles, as in A), in the absence of dynactin (red circles, as in D), or in the absence of an EB protein (black squares, as in E). Mean values from three separate experiments (each with approximately 50 kymographs) are shown; error bars are s. d. (**G**) Kymographs of GFP-dynein microtubule end tracking in the absence of ATP (in contrast to all other experiments that contain ATP). Other conditions as in A. For merged kymographs, microtubule plus and minus ends are labelled by (+) and (−). Experiments were performed at 30°C.

Using an Alexa647-labelled version of the SNAP-tagged EB1 homolog EB3 (Fig. EV1B) we observed that GFP-dynein indeed colocalised with the EB at growing microtubule ends in the presence of dynactin (Fig. 1C) in an ATP-independent manner (Fig. 1G). Omitting either dynactin (Fig. 1D) or EB (Fig. 1E) from the assay strongly reduced end tracking of GFP-dynein as evidenced by the averaged GFP-dynein intensity profiles at the plus ends. These results demonstrate that the human dynactin complex is sufficient to recruit the dynein complex to EBs at growing microtubule ends. Replacing human dynactin by purified neuronal pig brain dynactin (Fig. EV1B) resulted in similar microtubule end tracking behaviour of dynein (Fig. EV2), demonstrating that both purified dynactin complexes can form a link between EBs and dynein. Together, these results suggest that also in the context of the entire dynactin complex the CAP-Gly domain of p150 can be exposed, to allow end-tracking of dynein without the requirement of other regulators. These findings go beyond previous work with a p150 fragment and provide insight about the end tracking functionality of the entire dynactin complex (Duellberg et al., 2014).

The interaction between dynein and dynactin is stabilised by several cargo adaptors, such as, Bicaudal D2 (BicD2), which results in activation of processive dynein motion (Hoogenraad and Akhmanova, 2016; McKenney et al., 2014; Schlager et al., 2014). It is not known how this stabilisation and activation of processive motion influences the end tracking behaviour of dynein. To address this question, we purified an N-terminal fragment of the human cargo adaptor protein BicD2 (BicD2-N) (Fig. EV1B), which has been shown to trigger processive dynein-dynactin motility (McKenney et al., 2014; Schlager et al., 2014; Splinter et al., 2012). Addition of 200 nM Alexa647-labelled SNAP-tagged BicD2-N (Alexa647-BicD2-N) to a reconstitution assay with dynamic microtubules, dynein, dynactin and EB1 lead now to the appearance of processive and lattice-bound GFP-dyneins in addition to plus-end accumulated dynein (Fig. 2A). To our knowledge, this is the first observation of processive human dynein motility on dynamic microtubules using purified proteins.

**Figure 2.**
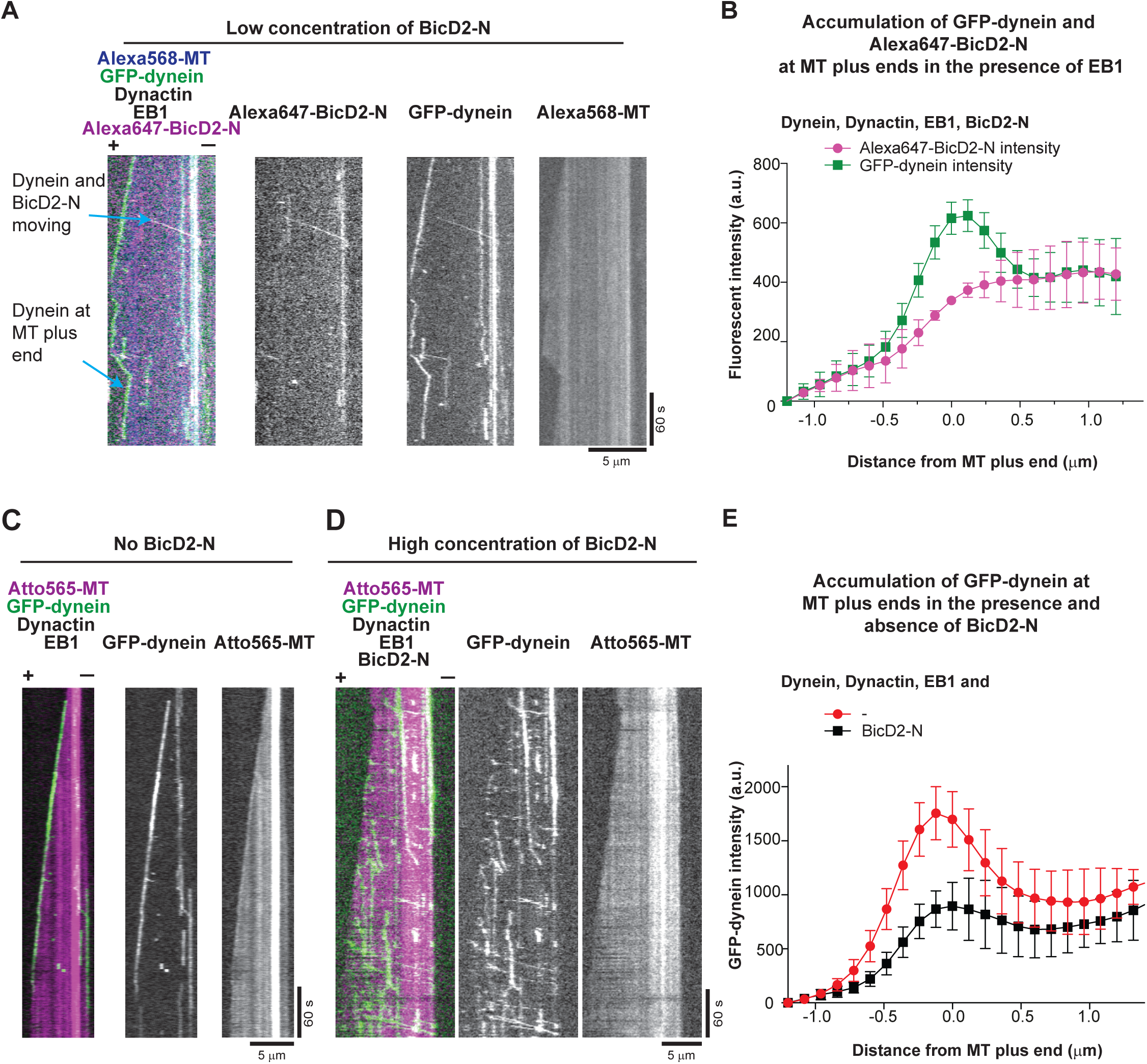
Effect of BicD2-N on microtubule end tracking of dynein in the presence of dynactin and EB1. (**A**) Merged triple colour and single fluorescence channel TIRF microscopy kymographs of GFP-dynein showing end tracking, processive motility and diffusive and static binding in the presence of all DDB components and EB1. Protein concentrations are 10 nM GFP-dynein (green in merge), 20 nM human dynactin, 20 nM EB1, 200 nM Alexa647-BicD2-N (magenta in merge), and 17.5 μM Alexa568-tubulin (blue in merge). Alexa647-BicD2-N often colocalises with processively moving and occasionally with statically bound GFP-dynein, but not with plus end-tracking GFP-dynein. (**B**) Averaged fluorescence intensity profiles of GFP-dynein (green circles) and Alexa647-BicD2-N (magenta squares) at growing microtubule plus ends. Mean values from three separate experiments (each with approximately 50 kymographs) are shown.; error bars are s. d. (**C, D**) Dual and single colour kymographs of an Atto565-microtubule (magenta) in the presence of GFP-dynein (green), human dynactin and EB1, and either **(C)** in the absence or **(D)** presence of 5 μM BicD2-N; other conditions as in A. The increased BicD2-N concentration in D compared to A strongly reduces microtubule plus end tracking of GFP-dynein. (**E**) Averaged fluorescence intensity profiles of GFP-dynein at growing microtubule plus ends for the condition without BicD2-N (black squares, as in C) or with 5 μM BicD2-N (red circles, as in D). Mean values from three separate experiments (each with approximately 50 kymographs) are shown; error bars are s. d. Microtubule plus and minus ends in merged kymographs are labelled by (+) and (−). Experiments were performed at 30°C.

BicD2-N colocalised mainly with processive GFP-dynein (Fig. 2A), as expected for the ternary complexes consisting of dynein, dynactin and BicD2-N (DDB) (McKenney et al., 2014; Schlager et al., 2014). Interestingly, BicD2-N did not colocalise with plus-end tracking GFP-dynein (Fig. 2A, B). Velocity and run length of processive DDB particles and the relative fractions of processive, diffusive and static DDB binding events on dynamic microtubules were similar to those reported for static microtubules (Fig. EV3) (McKenney et al., 2014; Schlager et al., 2014). Hence, the DDB complex was functional in our assay condition that supports microtubule dynamics. When reaching the dynamic microtubule minus ends, processive DDB particles accumulated there, also in the absence of EB1 (Fig. EV3).

Adding a large excess of BicD2-N (5 μΜ) to the end tracking experiment led to a strong reduction of EB1-dependent end tracking of GFP-dynein (Fig. 2C, D, Movie 2), as evidenced by averaged GFP-dynein intensity profiles at dynamic microtubule ends (Fig. 2E). At the same time, the increased concentration of BicD2-N induced more processive runs (Fig. 2D). These results show that DDB complex formation and processive motility are apparently incompatible with EB-mediated microtubule plus-end tracking of dynein.

This led us to ask whether this incompatibility is due to DDB particles processively leaving the EB-decorated end, which would predict that EBs would locally promote the initiation of runs from microtubule plus ends. To test this, we measured the relative initiation frequency of processive DDB runs as a function of the distance from the growing microtubule plus end (Fig. 3). Surprisingly, we observed a strongly increased initiation probability in the dynamic plus end region even in the absence of EB1 (Fig. 3A, B black bars, Fig. EV4 left). This strong bias was largely unaffected by the additional presence of EB1 (Fig. 3B red bars, Fig. EV4A right), which did also not strongly affect the intensity profile of GFP-dynein at growing microtubule plus ends in this condition (Fig. 3C). The strong bias of initiations of processive runs was specific for dynamic microtubule plus ends, as it was not observed for static, GMPCPP-stabilised microtubules, where only a much milder trend could be measured (Fig. 3D, Fig. EV4B). Together these data show that dynein/dynactin can exist as part of two different populations: either tracking plus ends in an EB-dependent manner or moving processively as part of a DDB complex. EB-dependent end tracking does not promote the initiation of dynein runs from plus ends, but instead DDB particles have a remarkable inherent preference for starting their runs from growing microtubule plus ends under the experimental conditions used in this study.

**Figure 3.**
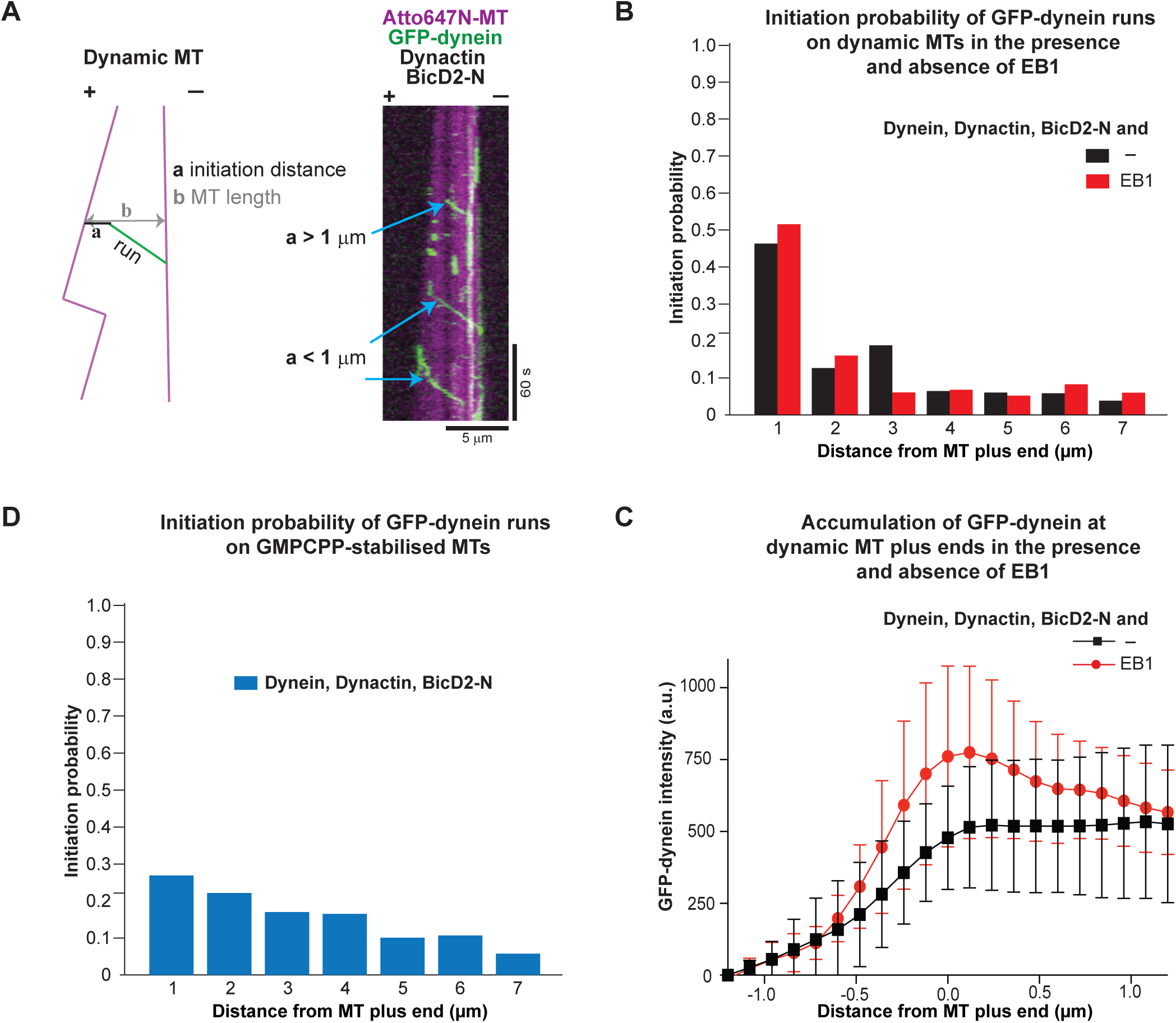
Spatial initiation probability of processive DDB runs. (**A**) Example dual colour kymograph and a schematic showing how the distance of initiation of processive runs was measured. All experiments contained 10 nM GFP-dynein, 20 nM pig dynactin, 5 μM BicD2-N. (**B**) Histograms of the spatial initiation probabilities of processive runs within the first 7 μm from growing microtubule plus ends in the absence (black bars), or presence of 10 nM EB1 (red bars). Alexa568-tubulin was present at 17.5 μM. (**C**) Averaged fluorescence intensity profiles of GFP-dynein at growing microtubule plus ends for the conditions as in B. Mean values from three separate experiments (each with approximately 50 kymographs) are shown; error bars are s. d. Experiments were performed at 30°C. (**D**) Histogram of spatial initiation probabilities of DDB on static GMPCPP-microtubules. Concentrations of DDB components as in B. Over 500 complexes were analysed for each conditions from three different data sets for B and C.

When investigating the reason for this bias of motility initiations, we noticed that increasing the dynactin concentration from 20 nM to 80 nM in our dynamic microtubule assay without EB proteins led to some visible plus end accumulation of GFP-dynein (Fig. 4A, Movie 3), suggesting that dynactin itself might be mediating direct microtubule plus end binding of the DDB complex. To probe this hypothesis further, we purified a fluorescent N-terminal fragment of the p150 subunit of dynactin (p150-N) (Fig. EV1B) and observed that 10 nM of p150-N could also recruit GFP-dynein to microtubule plus ends in the assay buffer used in this study, even in the absence of EB1 (Fig. 4B, Movie 4). Fluorescent p150-N clearly showed preferential localisation to growing microtubule ends, both in the presence (Fig. 4B and Movie 4) and absence (Fig. 4C) of dynein. Interestingly, p150-N bound also with high affinity to the stable GMPCPP segments of the microtubules used for surface immobilisation (Fig. 4C). These results demonstrate that in the assay buffer used here, the p150 dynactin subunit has an intrinsic, previously unreported preference for growing microtubule end localisation, mediated probably by recognising a conformational aspect of the GTP cap. This end accumulation however disappeared at higher ionic strength (Fig. 4D), explaining why in previous experiments, which were performed in higher ionic strength buffers, p150-N end accumulation was strictly dependent on EB1 (Duellberg et al., 2014), an interaction that appears to be less dependent on the ionic strength.

**Figure 4.**
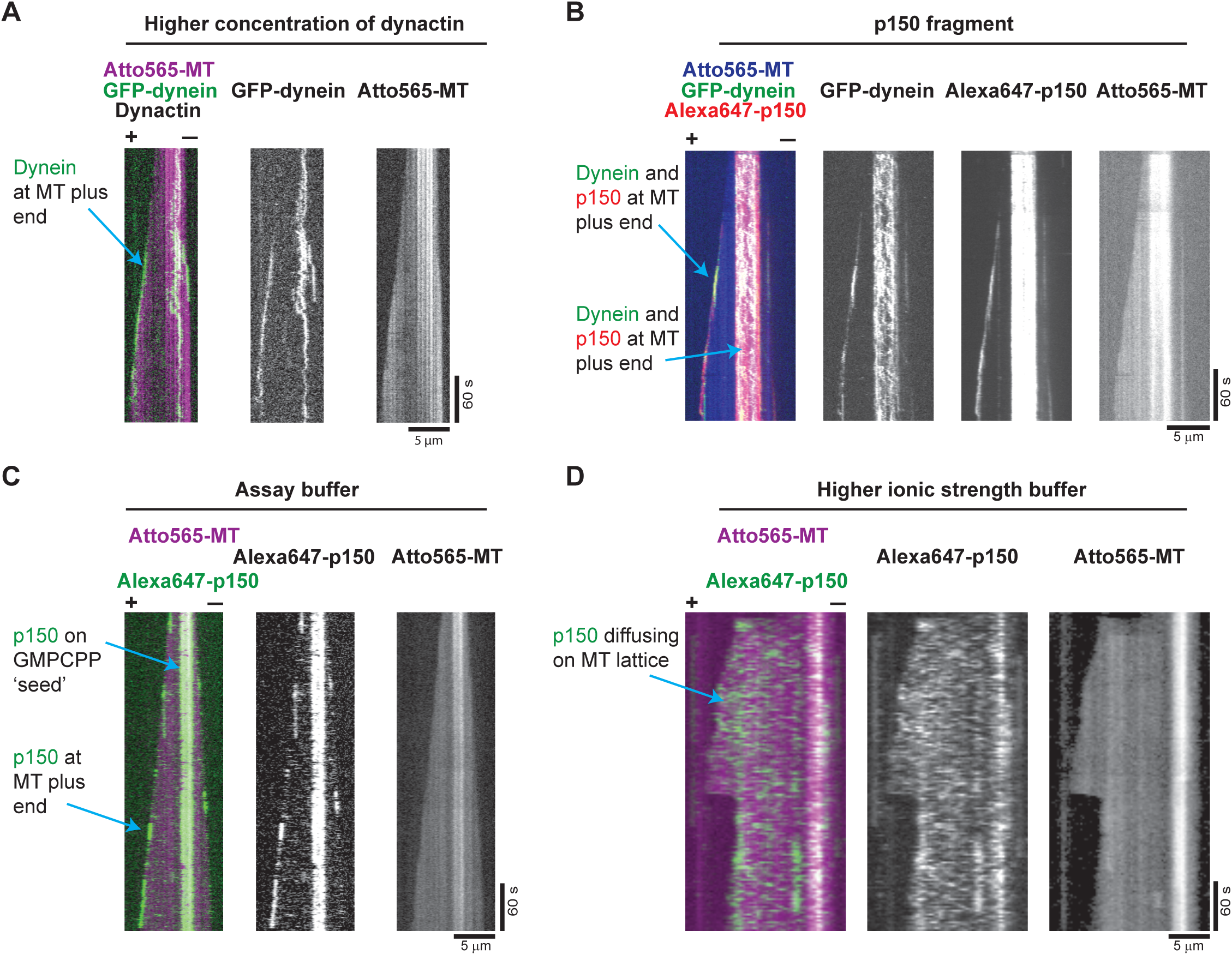
p150-dependent plus end tracking of dynein. **(A, B)** Multi and single colour kymographs showing under certain conditions p150 or dynactin-dependent microtubule plus end tracking of dynein in the absence of EB proteins: (**A**) 10 nM GFP-dynein (green) tracking the growing end of a Atto565-microtubule (magenta) in the presence of pig dynactin at an elevated concentration of 80 nM. (**B**) 10 nM GFP-dynein (green) tracking the plus end of an Atto565-microtubule (blue) in the presence of 10 nM Alexa647-p150 (a fragment containing the first 517 amino acids of p150) (red). Experiments in (A) and (B) were performed in standard assay buffer (BRB20 supplemented with 50 mM KCl, for details see Methods). (**C, D**) Dual and single colour kymographs showing that microtubule binding behaviour of p150 depends strongly on ionic strength: (**C**) In standard assay buffer, 10 nM Alexa647-p150 (green) accumulates at the plus end and on the GMPCPP-stabilised segment of a growing Atto565-microtubule (magenta). **(D)** At higher ionic strength (BRB80 supplemented with 60 mM KCl, see Methods), Alexa647-p150 (green) binds only weakly to the microtubule without a detectable plus end preference. Protein concentrations as in C. The Atto565-tubulin concentration was always 17.5 μM tubulin. The temperature was 30°C.

Taken together, our results demonstrate that in addition to the canonical EB1-dependent mechanism, there exists also a so far undescribed mechanism of dynein recruitment to growing microtubule ends depending on direct end recognition by the p150 subunit of the dynactin complex. This interaction explains the bias of the initiation of BicD2-dependent motility from microtubule plus ends, which is not further enhanced by EB proteins (see Discussion).

In metazoan cells, BicD2 and dynactin cooperate with yet another regulator, the Lis1 protein, to control both processive motion as well as end tracking of dynein (Splinter et al., 2012). To investigate the mechanism of the combinatorial interplay of these different dynein regulators, we purified a fluorescent Lis1 constructs (Fig. EV1B) and combined all purified regulators in our dynamic microtubule end tracking assay. Although 5 μM BicD2-N strongly reduced microtubule end tracking of dynein (Fig. 5A, and as shown earlier in Fig. 2D, Movie 2), combining 1 μM or 5 μM Lis1 with EB1, GFP-dynein, dynactin and BicD2-N showed that Lis1 can restore end tracking of GFP-dynein in a dose dependent manner (Fig. 5B, C, Movies 5, 6), as revealed by the GFP-dynein fluorescence profiles at growing microtubule plus ends (Fig. 5D). This activity of Lis1 can explain the requirement of Lis1 for end tracking of dynein in cells, where the presence of several cargo adaptors, including BicD2, may suppress EB-dependent end tracking of dynein (Coquelle et al., 2002; Splinter et al., 2012).

**Figure 5.**
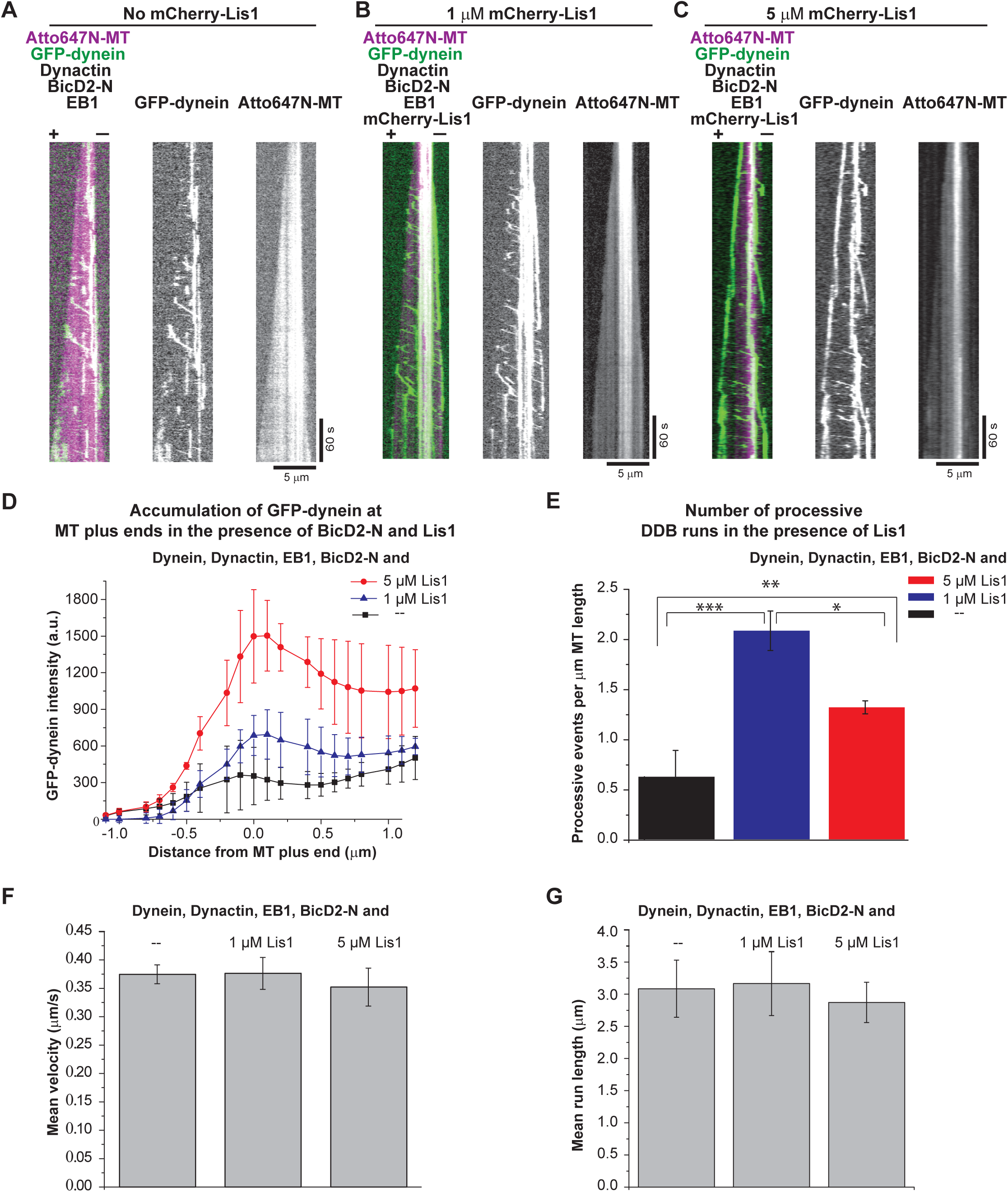
Regulation of microtubule end tracking versus processive motility of dynein by Lis1 and BicD2-N in the presence of dynactin and EB1. (**A-C**) Kymographs showing GFP-dynein (green) on dynamic Atto647N-microtubules (magenta) in the presence of all the regulators at different Lis1 concentrations. Protein concentrations were 10 nM GFP-dynein, 20 nM pig dynactin, 20 nM EB1, 5 μΜ BicD2-N, and (**A**) either no Lis1, (**B**) 1 μΜ mCherry-Lis1, or (**C**) 5 μΜ mCherry-Lis1. Together with BicD2, Lis1 increases the number of processive dynein runs and high concentrations of Lis1 restore plus-end localisation of dynein in the presence of BicD2-N. Microtubule orientation as indicated. (**D**) Averaged intensity profiles of GFP-dynein at growing microtubule plus-ends in the presence of EB1, dynactin and 5 μM BicD2-N (black squares, as in A), in the presence of 1 μM mCherry-Lis1 (blue triangles, as in B) and 5 μM mCherry-Lis1 (red circles, as in C). Mean values from three separate experiments (each with approximately 50 kymographs) are shown; error bars are s. d. (**E**) The average number of processive DDB runs per μm microtubule length for the three different Lis1 conditions (as in A, B and, C). Error bars are s. d. *p = 0.001, **p = 0.003, ***p = 0.001 (unpaired t-test). Over 500 complexes were analysed for each conditions from two different data sets. Bar graphs showing (**F**) the mean velocity and (**G**) the mean run length of processive DDB events in the presence of all proteins (as in A, B and C). Error bars are the s.e.m. Mean velocities were 0.37 ± 0.02 μm/s (A), 0.38 ± 0.03 μm/s (B), and 0.35 ± 0.03 μm/s (C). Mean run lengths were 3.1 ± 0.4 μm (A), 3.2 ± 0.5 μm (B), and 2.9 ± 0.3 μm (C). For each condition over 300 complexes were analysed from three different data sets. Experiments were performed at 30°C.

Remarkably, despite promoting microtubule end tracking of GFP-dynein, adding 1 μM of Lis1 together with all the purified regulators also strongly increased the number of processive dynein runs (Fig. 5E). However, contrary to the previously reported decrease in the velocity of dynein in gliding assays (Torisawa et al., 2011; Wang et al., 2013; Yamada et al., 2008), we found that neither the velocity nor the run length of the processive DDB complexes moving on dynamic microtubules were affected by the presence of Lis1 (Fig. 5F, G, respectively). For most of the DDB particles we were also unable to visualise fluorescently labelled Lis1 comigrating with processive GFP-dynein in triple colour imaging experiments (Fig. EV5), suggesting that Lis1 either did not bind processive DDB complexes or, alternatively, Lis1 does not affect their velocity and run length. Interestingly, triple colour imaging of Lis1 in the end tracking assay with all the regulators clearly showed some accumulation of Lis1 at growing microtubule plus ends (Fig. EV6A). In addition, the initiation probability of processive GFP-dynein runs from the microtubule plus end region also increased further in the additional presence of Lis1 (Fig. EV6B, C). Together these data suggest that in the presence of all dynein regulators studied here, Lis1 has a dual role. First, it can act as an initiation factor for processive dynein motility for which BicD2-N is additionally required. Lis1 can contribute to enhance both the efficiency of initiation as well as its plus end bias. Second, Lis1 can also promote recruitment of dynein/dynactin to growing microtubule plus ends by binding to dynein/dynactin without BicD2-N, thereby depleting the pool of motion-competent DDB particles at high Lis1 concentrations.

## DISCUSSION

In this study, we have dissected the molecular mechanism of how three major dynein regulators control the balance between microtubule end tracking and processive motility of human dynein. Our results reveal that dynactin has a triple role: (i) it mediates EB-dependent plus end tracking of dynein, in the absence of other regulators, (ii) it activates BicD2-dependent dynein processivity, and (iii) it establishes a bias in dynein motility initiation from microtubule plus end regions by also interacting with microtubule plus end regions autonomously. Remarkably, EB-dependent end tracking of dynein/dynactin does not contribute to biasing motility initiation towards microtubule plus ends. The microtubule recruitment factor Lis1 adds an additional layer of regulation. It is required for efficient EB-dependent plus-end tracking of dynein/dynactin in the presence of BicD2; and in cooperation with BicD2, Lis1 can also promote initiation of processive motility.

Recent structural data suggested a possibility of dynactin adopting an autoinhibited state with its p150 CAP-Gly domain and the first coiled-coil being inaccessible for interaction with EBs and the dynein intermediate chain (Urnavicius et al., 2015). This raised the question of how relevant was the previous reconstitution with an N-terminal fragment of p150 mediating EB1-dependent end tracking of dynein (Duellberg et al., 2014). Here we found that the p150 CAP-Gly domain in the context of the entire dynactin complex is free to associate with EBs in agreement with the variable orientations of the p150 N-terminus that were also observed by cryo-electron microscopy (Urnavicius et al., 2015). The previously noticed weak and likely transient interaction between the dynein and dynactin complexes (McKenney et al., 2014) is apparently sufficient to support EB-dependent microtubule end tracking (Duellberg et al., 2014; Moughamian and Holzbaur, 2012; Moughamian et al., 2013; Peris et al., 2006), as our reconstitution with the entire dynactin complex shows here.

In our in vitro reconstitutions we also observed dynein tracking shrinking microtubule ends. This behaviour has also been observed in cells (Ten Hoopen et al., 2012) and in vitro with microsphere-immobilised dynein in the absence of dynactin (Hendricks et al., 2012; Laan et al., 2012). Future experiments with simultaneously fluorescently labelled individual dynein and dynactin molecules will be required to elucidate the mechanism of shrinking end tracking of dynein and dynactin.

Our combined observations that the dynactin component p150 has the previously undescribed property of strongly biasing processive dynein motility initiation towards microtubule plus end regions and that EB proteins do not regulate this bias predict that processive DDB complexes may not interact with EB proteins. This would indicate that in the DDB complex, the p150 subunit of dynactin might be conformationally restricted so that its affinity for binding the microtubule surface might be considerably higher than for binding the C-terminus of EB proteins. Alternatively, EBs might block the p150 binding site on microtubules. However, since BicD2-N was never observed to track growing microtubule ends in the presence of end tracking dynein/dynactin, we consider it more likely that the DDB complex is in a conformational state that is less likely to interact with EB1 than with the microtubule directly. This scenario suggests a potentially simple competition-based explanation for why overexpression of the BicD2-N fragment in human cells abolished plus end tracking of dynein (Splinter et al., 2012). A similar mechanism for down-regulating dynein end tracking by adapter proteins appears to exist in budding yeast where the expression of the cortical adaptor Num1 also removed dynein from microtubule plus ends (Lammers and Markus, 2015).

The intrinsic bias for processive motility initiation of DDB complexes may be explained by a selective recognition of the GTP cap region of growing microtubule plus ends by p150, especially since p150 was also observed to bind preferentially to GMPCPP microtubules. This property is strikingly similar to that of the unrelated microtubule binding protein TPX2 that was recently also observed to strongly bind to GMPCPP microtubules and to growing microtubule ends independently of EB proteins (Roostalu et al., 2015). Possibly, this property is related to the reported microtubule nucleation and microtubule catastrophe suppressing activity of p150 and TPX2 (Lazarus et al., 2013; Roostalu et al., 2015). In the future, it will be interesting to study how p150 interacts with the microtubule in its GTP-bound state as this is likely the relevant configuration for transport initiation. It will be interesting to understand how this interaction mode compares to the reported structure of the p150 CAP-Gly domain and its neighbouring acidic region binding in a flexible manner to the C-terminal tubulin tails of GDP/taxol microtubules (Wang et al., 2014).

In cultured mammalian cells, end tracking of dynein was reported to be reduced in the absence of Lis1 (Coquelle et al., 2002; Splinter et al., 2012). This can be explained by our finding that Lis1 appears to compete with the adapter protein BicD2 counteracting the BicD2-mediated reduction of dynein/dynactin end tracking. This suggests that in cells the negative effect of cargo adapters on dynein end tracking would be, at least in part, compensated for by Lis1. This is in agreement with data from budding yeast where overexpression of the Lis1 homologue Pac1 rescued plus end localisation of dynein in cells overexpressing Num1 adaptor protein (Lammers and Markus, 2015).

Several biochemical and structural data hint at the possibility of an allosteric mechanism underlying the competition between Lis1 and BicD2 for dynein binding. Although both regulators are known to interact with both dynein and dynactin, currently no overlapping binding sites are known. Lis1 binds the AAA+ ring of the dynein motor domain (Huang et al., 2012; Tai et al., 2002) and the p50 subunit of dynactin (Tai et al., 2002), whereas BicD2 interacts with the dynein tail and the Arp1 filament of dynactin (Chowdhury et al., 2015; Urnavicius et al., 2015). Interestingly, Lis1 has been shown to bind to a compact dynein conformation with docked motor domains (also known as the phi particle) (Toba et al., 2015) that has been proposed to correspond to an autoinhibited state of the dynein motor (Torisawa et al., 2014). In contrast, in the DDB particle the motor domains of dynein show a splayed configuration, possibly corresponding to its processive conformation (Carter et al., 2016; Urnavicius et al., 2015). Therefore, it is possible that Lis1 and BicD2 bind preferentially to different dynein conformations, and their competition might shift the equilibrium between the inactive and processive dynein configurations. As such, our data also predict that the EB and dynactin-dependent plus end tracking dynein particles may still be in a compact or inhibited state as splaying of dynein would additionally require interaction with BicD2.

Lis1 is well known to be involved in the initiation of dynein-dependent transport in cells (Egan et al., 2012; Lenz et al., 2006; Moughamian et al., 2013). Reducing Lis1 levels in fungi and mammalian cells inhibits dynein-dependent transport of several organelles (Dix et al., 2013; Egan et al., 2012; Klinman and Holzbaur, 2015; Moughamian et al., 2013; Pandey and Smith, 2011) and BicD2-N carrying vesicles (Splinter et al., 2012) in agreement with the simultaneous requirement of Lis1 and BicD2 for transport initiation as observed in our minimal system. At the same time, reduction of Lis1 levels in Drosophila cells was recently shown to enhance dynein-dependent transport of mitochondria (Vagnoni et al., 2016), suggesting that Lis1 can also have an inhibitory effect on transport initiation, potentially when it is in excess over cargo adaptors. The contrasting behaviours observed in cells may be explained by the possible existence of an optimal Lis1/adapter protein ratio for efficient initiation of processive motility, as our in vitro experiments suggest.

Reports of the effect of Lis1 on the velocity of dynein motility vary. In vitro experiments with purified proteins showed that Lis1 slowed down dynein motility in gliding assays where teams of surface-immobilised motors propel microtubules (Torisawa et al., 2011; Wang et al., 2013; Yamada et al., 2008). The presence of dynactin reverted this slow-down (Wang et al., 2013). In filamentous fungi Lis1 (Egan et al., 2012) or in *in vitro* single molecule experiments with purified proteins in the absence of mechanical load (McKenney et al., 2010), the velocity of dynein was unaffected by the presence of Lis1, similar to our observation of Lis1 not affecting the velocity of dynein. It therefore appears that at least under unloaded conditions Lis1 may be considered a microtubule initiation factor that may release from DDB when processive motility is activated.

In conclusion, this study establishes a separate role for each of the three key regulators of dynein activity and shows that dynein can exist in two functionally different pools, either as a non-motile EB-dependent microtubule plus end-tracking complex or as part of a DDB complex with activated processivity. In the DDB complex, the CAP-Gly domain of the dynactin subunit p150 is responsible for biasing initiation of processive motility at microtubule plus ends and on freshly grown tyrosinated microtubules (McKenney et al., 2016). Lis1 acts as a general initiation factor that appears to compete in a complex manner with processivity enhancing adapter proteins for dynein and dynactin binding. Biochemical separation of these two dynein pools might allow for independent control of dynein delivery to target structures and of dynein-mediated processive transport.

In the future, structural studies of the dynein complex together with its three major regulators will likely provide important mechanistic insight into the conformations of dynein and dynactin during plus-end tracking versus initiation of transport. In parallel, further extensions of dynamic in vitro reconstitutions promise to lead to the dissection and mechanistic understanding of additional layers of regulation of dynein activity.

## MATERIALS AND METHODS

### Plasmids

To generate a Multibac expression construct of the human cytoplasmic dynein complex labelled with mGFP, we first replaced the SNAP-tag sequence in the pACEBac1 vector (Vijayachandran et al., 2013) (generous gift from A. Carter) with the mGFP sequence to generate a plasmid (pGFPdyn1) encoding for a fusion protein consisting of an N-terminal His_6_-tag followed by a ZZ-tag, a TEV protease cleavage sequence, mGFP and the human cytoplasmic dynein heavy chain (His_6_-ZZ-TEV-mGFP-DHC). As described (Schlager et al., 2014), using purified Cre recombinase (New England Biolabs) this plasmid was combined with pIDC (Vijayachandran et al., 2013) (generous gift from A. Carter) that contained the accessory human dynein subunits IC2C, LIC2, Tctex1, LC8 and Robl1 (pdyn2), producing a construct for the simultaneous expression of all six dynein subunits in insect cells. The presence of all subunits in the recombined construct (pGFPdyn3) was confirmed by PCR.

The coding sequence of full-length human Lis1 was amplified from a cDNA (accession number BC064638) by PCR and was cloned into a pFastBac1 plasmid (Invitrogen) to generate a construct for insect cell expression of an N-terminally tagged His_6_-mCherry-Lis1 or His_6_-SNAP-Lis1 where the His_6_ tag was separated from the mCherry or SNAP sequence by a TEV protease cleavage site.

The coding sequence for the N-terminal 400 amino acids of human BicD2 was amplified from a cDNA (Origen, SC300552) by PCR and cloned into a pETZT2 plasmid. Sequences were added to generate a bacterial expression construct encoding for a His_6_-tag followed by a Z-tag, a TEV cleavage site, a SNAP tag, a Gly5 linker, and the N-terminal BicD2 fragment (His_6_-SNAP-BicD2-N in brief).

The coding sequence of full-length human EB3 was amplified by PCR from the plasmid pET28a-His-mCherry-EB3 (generous gift from M. Steinmetz), keeping the ‘long linker’ between mCherry and EB3. Sequences were combined in a pETMZ plasmid to generate a bacterial expression construct encoding for a fusion protein consisting of a N-terminal His_6_-tag followed by a Z-tag, a TEV protease cleavage site, a SNAP tag, the ‘long linker’, and EB3 (His_6_-SNAP-EB3 in brief).

The sequences of all constructs were verified by sequencing.

### Purification of recombinant human dynein

Human GFP-dynein was expressed in *Sf*21 insect cells and purified by immobilized metal affinity chromatography (IMAC) followed by size exclusion chromatography, as previously described (Trokter et al., 2012). All purification steps were carried out at 4°C and the buffers were degassed and chilled to 4°C. Briefly, a pellet of 800 ml of cell culture was thawed on ice and resuspended in 100 ml lysis buffer (50 mM HEPES pH 7.4, 250 mM K-acetate, 20 mM imidazole 2 mM MgSO_4_, 0.25 mM EDTA, 10% glycerol (vol/vol), 0.2 mM Mg-ATP, 1 mM β-mercaptoethanol (ßME)) supplemented with protease inhibitors (Complete EDTA Free, Roche Applied Science). Cells were lysed using a dounce homogenizer (Wheaton). After clarification of the lysate by ultracentrifugation (183,960 × *g*, 45 min, 4°C), the supernatant was loaded onto a 5 ml HiTrap chelating column (GE Healthcare) loaded with CoCl_2_. The column was then washed with 200 ml lysis buffer. The protein was eluted using elution buffer (50 mM HEPES, pH 7.4, 250 mM K-acetate, 350 mM imidazole, 2 mM MgSO_4_, 0.25 mM EDTA, 10% glycerol (vol/vol), 0.2 mM Mg-ATP, 1 mM ME). The dynein containing fractions were pooled and exchanged to gel filtration buffer (30 mM HEPES pH 7.4, 150 mM K-acetate, 2 mM MgSO_4_, 0.5 mM EGTA, 10% glycerol (vol/vol), 0.05 mM Mg-ATP, 10 mM ßME). The His_6_ tag was cleaved off by incubating with a TEV protease overnight at 4°C. After TEV cleavage, the protein was concentrated to a 5 ml volume using Vivaspin concentrator (Sartorius) with a 50 kDa molecular weight cut-off. The concentrated protein was further purified by size-exclusion chromatography using a Superose 6 XK 16/70 prep grade column (GE Healthcare) equilibrated in gel filtration buffer. The mGFP-dynein complex containing peak fractions were identified by SDS-PAGE Coomassie Blue G-250 staining. The fractions of interest were pooled and concentrated to approximately 0.3 mg/ml using a Vivaspin 50 kDa molecular weight cut-off concentrator and ultracentrifuged (174,000 × *g*, 10 min, 4°C). Protein aliquots were flash frozen and stored in liquid nitrogen.

### Dynactin purification from HeLa S3 cells

Native human dynactin complex was purified from HeLa S3 cells by BicD2-N affinity chromatography followed by ion exchange chromatography. To generate a BicD2-N column for dynactin affinity purification, 60 mg of purified His_6_-SNAP-BicD2-N (see below) was conjugated to a 5 ml NHS column (GE Healthcare) following the conjugation method described by (Widlund et al., 2012).

Frozen cell pellets from 25 l HeLa S3 cell culture were thawed and resuspended in approximately 150 ml lysis buffer (30 mM HEPES, pH 7.2, 50 mM K-acetate, 1 mM EGTA, 5 mM Mg-acetate, 0.5 mM ATP, 2 mM ME), supplemented with 10 μg/ml DNAse I, and protease inhibitors (Complete-EDTA Free, Roche Applied Science) using a Polytron tissue dispenser by three pulses of 6.0 ×10^3^ rpm for 90 sec interspersed by 90 s incubations on ice. The lysate was clarified by centrifugation (125,200 × g, 40 min, 4°C). The supernatant was recovered and filtered using a 0.22 μm Steritop filter (Millipore, Bedford, MA), and loaded onto a BicD2 column at 0.5 ml/min. The column was washed with 10 column volumes of wash buffer (lysis buffer supplemented with 100 mM NaCl). The dynein-dynactin complex was then eluted by applying a linear NaCl gradient in lysis buffer using an Akta Purifier FPLC system (GE Healthcare). The dynactin enriched fractions were identified by SDS-PAGE analysis. Four such BicD2 affinity purifications were run in parallel per day (4 BicD2 columns, 4 x 25 l HeLa S3 cell lysate), for a total of three days (300 l HeLa S3 cell lysate in total). The dynactin containing fractions from all rounds of BicD2 affinity purification were pooled, flash frozen and stored at -80°C. Between each round of purification the four BicD2 columns were extensively washed with 1 M NaCl and 500 mM NaCl dissolved in lysis buffer and stored at 4°C. For long term storage, the BicD2 columns were washed with 100 ml of 6x PBS, then with 100 ml of 1x PBS and finally with 1x PBS with 50% glycerol for storage at -20°C.

The pooled dynactin enriched fractions were thawed followed by buffer exchange to MonoQ buffer (35 mM Tris pH 7.2, 5 mM MgSO_4_, with 0.5 mM ATP and 2 mM ME) using HiPrep 26/60 desalting columns (GE Healthcare). After buffer exchange the protein was ultracentrifuged (174,000 × g, 20 min, 4°C) and loaded on a MonoQ 5/50 GL column (GE Healthcare) pre-equilibrated with MonoQ buffer. The dynactin complex was eluted by applying a step gradient of NaCl dissolved in MonoQ buffer using the elution regime described by (Bingham et al., 1998). The dynactin complex eluted at 390 mM NaCl as identified by Coomassie Blue stained SDS-PAGE analysis, and further verified by western blotting using an anti-p150 antibody (BD-bioscience) and mass-spectrometry. The dynactin fractions were pooled and buffer exchanged to storage buffer (30 mM HEPES, pH 7.2, 50 mM K-acetate, 1 mM EGTA, 2 mM Mg-acetate, 0.5 mM ATP, 2 mM ME, 10% glycerol), concentrated to 0.3 mg/ml using Vivaspin concentrator (Sartorius) with a 50 kDa molecular weight cut-off, ultracentrifuged (174,000 ×g, 10 min, 4°C), flash frozen in 5 μl aliquots and stored in liquid nitrogen. Approximately 0.8 mg of human dynactin was obtained from 300 l HeLa S3 cell culture. The final protein concentration was 0.3 mg/ml.

### Dynactin purification from pig brain

Native dynactin was also purified from pig brain following the same method as described above. Compared to the large volume of starting material required for purification of human dynactin from HeLa S3, a much smaller volume of material is required when purifying from pig brains, offering a faster alternative for the purification of mammalian dynactin. Compared to the purification from Hela S3 cells, the purification from pig brains differed in the following manner: (1) Two frozen pig brains were cut into small pieces with a scalpel and supplemented with 2 volumes of ice-cold lysis buffer. The brains were then thawed on ice and homogenized using a Polytron dispenser following the same cycle as described for the HeLa S3 cell lysis above. (2) The homogenate after lysis was centrifuged twice, first: 29,000 × g, 30 min, at 4°C; and second: 125,200 × g, 40 min, 4°C. (3) The clarified, unfiltered lysate from two brains was loaded onto one BicD2 column, and the first round of purification was done in one day. The BicD2 column eluate was stored at 4°C overnight and loaded onto a MonoQ 5/50 HR column the next day, following the same procedure as described for purification from the Hela S3 cells. Approximately 0.2 mg of dynactin was obtained from two pig brains. The final protein concentration was 0.2 mg/ml. The experiments with Lis1 (Fig. 5, and Fig. EV5, 6) were performed using purified pig dynactin.

### Purification and labelling of BicD2-N

His_6_-SNAP-BicD2-N was expressed in *E.coli* (BL21 pRIL) by induction with 1 mM IPTG for 16 hours at 18°C and purified by IMAC as described below. The pellets from 4 liters of cell culture were thawed and resuspended in lysis buffer (50 mM HEPES pH 7.4, 100 mM KCl, 10 mM imidazole, 1 mM ME, 0.1 mM ATP), supplemented with protease inhibitors (Complete-EDTA Free, Roche Applied Science). The cells were lysed using a microfluidizer. The lysate was clarified by ultracentrifugation (184,000 × *g*, 45 min, 4°C) and loaded onto a 5 ml HisTrap column (GE Healthcare) pre-equilibrated with the lysis buffer. The column was then extensively washed with lysis buffer. The protein was eluted with elution buffer (50 mM HEPES pH 7.4, 100 mM KCl, 300 mM imidazole, 1 mM ME, 0.1 mM ATP). The protein containing fractions were pooled and dialysed into gel filtration buffer (50 mM HEPES, pH 7.4, 100 mM KCl, 1 mM ßME, 0.05 mM ATP, 10% (vol/vol) glycerol). The His_6_ tag was then cleaved off by overnight incubation with TEV protease at 4°C. After TEV cleavage, the SNAP-BicD2-N protein was concentrated and further purified by gel filtration using a Superdex 200 10/300 GL column (GE Healthcare) pre-equilibrated with gel filtration buffer. The fractions containing SNAP-BicD2-N (BicD2-N in brief) were pooled, concentrated to 2 mg/ml, ultracentrifuged (174,000 × *g*, 10 min, 4°C), and flash frozen in 5 μl aliquots in liquid nitrogen.

To generate a fluorescently labelled SNAP-BicD2-N (referred to as Alexa647-BicD2-N), the His_6_-SNAP-BicD2-N was incubated overnight at 4°C with an equimolar concentration of SNAP-Surface Alexa Fluor 647 (New England Biolabs) during the TEV protease cleavage reaction. The labelled protein was then passed through HiPrep 26/60 desalting columns (GE Healthcare) to remove the unreacted dye. Size exclusion chromatography and protein flash freezing was performed as described for the unlabelled protein. Final labelling ratio was 0.81 fluorophores per BicD2-N (monomer).

The SNAP-BicD2-N protein that was used for generating the BicD2 column for dynactin purification was dialysed into gel filtration buffer after IMAC elution, ultracentrifuged (174,000 × *g*, 10 min, 4°C), flash frozen and stored at -80°C. The final protein concentration was here 0.9 mg/ml.

### Purification of EB1 and EB3

Untagged EB1 was expressed and purified as described (Roostalu et al., 2015). The final protein concentration was 1.2 mg/ml. To generate Alexa647-EB3, His_6_-SNAP-EB3 was expressed in *E.coli* (BL21 pRIL) induced by 0.2 mM IPTG for 16 hours at 18°C and purified by IMAC as follows The pellets were thawed and resuspended in lysis buffer (50 mM NaPi pH 7.2, 400 mM KCl, 5 mM MgCl_2_, 0.5 mM ME) supplemented with protease inhibitors (Complete-EDTA Free, Roche Applied Science). The lysate was clarified by centrifugation (184,000 × g, 45 min, 4°C) and passed over a 5 ml HiTrap Chelating column (GE Healthcare) loaded with CoCl_2_ pre-equilibrated with lysis buffer. The column was then extensively washed with wash buffer (50 mM NaPi, pH 7.2, 400 mM KCl, 15 mM imidazole, 5 mM MgCl_2_, 0.5 mM ME). The protein was eluted with elution buffer (50 mM NaPi, pH 7.2, 400 mM KCl, 400 mM imidazole, 5 mM MgCl_2_, 0.5 mM ME). The protein containing fractions were pooled and the buffer was changed to lysis buffer using PD-10 Desalting columns (GE Healthcare). In order to fluorescently label the protein, an equimolar concentration of SNAP-Surface Alexa Fluor 647 (NEB) was added followed by overnight incubation at 4°C in the simultaneous presence of TEV protease cleavage. The unreacted dye was then removed by buffer exchange via HiPrep 26/60 desalting columns (GE Healthcare) followed by size exclusion chromatography using a Superdex 200 10/300 GL column (GE Healthcare). The peak fractions containing Alexa647-labelled SNAP-EB3 were pooled, ultracentrifuged (174,000 × *g*, 10 min, 4°C), aliquoted and flash frozen in liquid nitrogen. The final protein concentration was 0.6 mg/ml

### Purification of Lis1 constructs and labelling

His_6-_-mCherry-Lis1 was expressed in Sf21 cells. The cell pellet from a 600 ml culture was resuspended in lysis buffer (50 mM HEPES, pH 7.4, 100 mM KCl, 1 mM ME, 0.05 mM ATP, 10% (vol/vol) glycerol), supplemented with protease inhibitors (Complete-EDTA Free, Roche Applied Science). The cells were lysed using a dounce homogenizer. The lysate was first clarified by centrifugation (184,000 × *g*, 45 min, 4°C) followed by incubation with 1 g of Proteino Ni-TED resin (Macherey-Nagel) in a batch format. The resin was then transferred to a column and extensively washed with lysis buffer. The bound His_6_-mCherry-Lis1 was cleaved and released from the resin by TEV protease dissolved in lysis buffer incubated for 1 h at room temperature while re-suspending the resin every 5 min. The resin was then separated from the eluted protein by centrifugation (at 1,000 × *g*, 2 min, 4°C). The mCherry-Lis1 protein was further purified by gel filtration using a Superdex 200 10/300 GL column (GE healthcare) preequilibrated with lysis buffer. The mCherry-Lis1 containing fractions were identified by SDS-PAGE page, pooled, ultracentrifuged (174,000 × *g*, 10 min, 4°C), supplemented with 20% glycerol (vol/vol), flash frozen in 5 μl aliquots, and stored in liquid nitrogen. The final protein concentration was 2 mg/ml.

To generate a fluorescently labelled SNAP-Lis1 (referred to as Alexa647-Lis1), the His_6_-SNAP-Lis1 was expressed in Sf21 cells and purified as described for mCherry-Lis1 with the following modification: after the first round of gel filtration, purified SNAP-Lis1 was incubated overnight at 4°C with an equimolar concentration of SNAP-Surface Alexa Fluor 647 (New England Biolabs). The labelled protein was then passed through a HiPrep 26/60 desalting columns (GE Healthcare) to remove the unreacted fluorophore, followed by a second round of gel filtration using Superdex 200 10/300 GL column (GE healthcare). The purified Alexa647-Lis1 was flash frozen at a final protein concentration of 0.6 mg/ml.

### Purification and labelling of p150

His_6_-SNAP-p150-N (containing the first N-terminal 547 amino acids of the neuronal isoform of p150^Glued^) was purified and labelled with Alexa647 as described (Duellberg et al., 2014).

### Tubulin purification and labelling

Porcine brain tubulin was purified as described in (Castoldi and Popov, 2003) and covalently labelled either with NHS-biotin (Pierce), NHS-Alexa568 (Life technology), NHS-Atto565 (Sigma Aldrich) or NHS-Atto647N (Sigma Aldrich) as described by (Hyman, 1991).

### Total internal reflection fluorescence (TIRF) microscopy

The flow cells for TIRF microscopy experiments were assembled from a passivated biotin-PEG (polyethylene glycol) functionalized glass coverslips and a poly (L-lysine)-PEG (SuSoS)-passivated counter glass as described previously (Bieling et al., 2010). Fluorescently labelled biotinylated GMPCPP-stabilized microtubule ‘seeds’ for TIRF assays with dynamic microtubules were prepared as described previously (Bieling et al., 2010) (containing 10% of either Alexa568-, Atto565-. or Atto647N- labelled tubulin). The assay was modified from the protocol developed earlier (Bieling et al., 2010). Briefly, a flow cell was first incubated for 5 min with 5% Pluronic F-127 (Sigma-Aldrich) at room temperature, washed with BRB80 (80mM PIPES, 1mM EGTA, 1mM MgCl_2_) supplemented with 50 μg/ml κ-casein (BRB80κ) followed by an incubation with 50 μg/ml of NeutrAvidin in BRB80κ (Life Technologies) on ice for 5 min and washed with BRB80. After warming the flow cell to room temperature, Alexa568-labelled biotinylated GMPCPP ‘seeds’ diluted in BRB80 were passed through and incubated for 5 min for attachment, and then sequentially washed with BRB80, followed by Assay Buffer (AB) (20 mM K-PIPES (pH 6.9), 50 mM KCl, 1 mM EGTA, 2 mM MgCl_2_, 5 mM ME, 0.1% methylcellulose (4,000 cp, Sigma), 20 mM glucose, 2 mM GTP and 2 mM ATP). The final assay mixture was then passed through the flow cell which was then sealed using vacuum grease (Beckman), immediately followed by TIRF microscopy imaging.

The final assay mixtures for the different experiments always consisted of (i) a concentrated protein mix of different stored proteins that was first incubated for 5 min (except DDB which was incubated for 10 min) at 4°C, was then diluted by the addition of (ii) 6 μl oxygen scavenger - tubulin mix (5× 180 μg/ml catalase (Sigma), 5× 750 μg/ml glucose oxidase (Serva), 5× 17.5 μM of tubulin, containing 4% of either Alexa568-, Atto565-, or Atto647N-tubulin, in BRB80), and was finally brought to 30 μl by adding (iii) the appropriate volume of AB. The final tubulin concentration was always 17.5 μM. For the different experiments, the concentrated protein mixtures and the final protein concentrations were:

#### GFP-dynein/dynactin/BicD2-N (DDB) motility assay on dynamic microtubules

Concentrated protein mix A (or A’) consisted of dynein, human dynactin and BicD2-N (or Alexa647-labelled SNAP-BicD2-N) at a molar ratio of 1:2:20 in 3.6 μl. Final protein concentrations in the assay were 10 nM GFP-dynein, 20 nM dynactin and 200 nM BicD2-N. For control experiments in the absence of dynactin or BicD2-N, their respective storage buffers were added instead of the protein to maintain the same buffer composition.

#### GFP-dynein microtubule end tracking assay

Concentrated protein mix B consisted of GFP-dynein, human dynactin and EB1 (diluted 1:100 in AB) at a molar ratio of 1:2:2 in 4.8 μl. The final protein concentrations in the assay were 10 nM GFP-dynein, 20 nM human dynactin, 20 nM EB1. For triple colour imaging of microtubule end tracking (Fig. 1C), the concentrated protein mix B’ contained Alexa647-labelled SNAP-EB3 instead of untagged EB1 in 3.1 μl, keeping the molar ratios the same and yielding the same protein concentrations in the assay. For control experiments without dynactin, dynactin storage buffer was added instead of dynactin. For experiments in the absence of ATP (Fig. 1G), ATP was omitted from AB, and GFP-dynein and human dynactin were exchanged to the AB without ATP.

#### Simultaneous GFP-dynein motility and end tracking assay

For triple colour-imaging (Fig. 2A), concentrated protein mix D contained GFP-dynein, human dynactin, EB1, and Alexa647-BicD2-N at a ratio of 1:2:2:20 in 6 μl. For dual colour imaging (Fig. 2D and 5A), concentrated protein mix E contained untagged SNAP-BicD2-N (buffer exchanged to AB) instead of labelled BicD2-N and the protein ratio was changed to 1:2:2:500 in 9.8 μl. Final protein concentrations in the assay were 10 nM GFP-dynein, 20 nM human dynactin, 20 nM EB1, and 200 nM Alexa647-BicD2-N or 5 μΜ BicD2-N.

For experiments with Lis1 (Fig. 5B, C and Fig. EV5, 6), concentrated protein mix F or F’ contained GFP-dynein, pig dynactin, EB1, BicD2-N (buffer exchanged to AB), and mCherry-Lis1 at a ratio of 1:2:2:500:100 or 1:2:2:500: in 14.9 μl. Final protein concentrations were 10 nM GFP-dynein, 20 nM pig dynactin, 20 nM EB1, 5 μM BicD2-N, 1μM or 5 μM mCherry-Lis1. For experiments without Lis1 or with lower mCherry-Lis1 concentrations, the buffer composition was kept constant by adding mCherry-Lis storage buffer instead of mCherry-Lis1.

### TIRF Imaging

All samples were imaged at 30°C on a TIRF microscope (iMIC, FEI Munich) described in detail previously (Duellberg et al., 2014; Maurer et al., 2014; Roostalu et al., 2015). Image acquisition and channel alignment were carried out as explained previously (Maurer et al., 2014). All time-lapse movies were recorded at 1 fps for 500 s. The exposure time was always 200 ms for all the channels. For dual colour TIRF microscopy imaging, GFP-dynein and Alexa568- or Atto565-labelled microtubules were simultaneously excited at 488 nm and 561 nm respectively.

For triple colour imaging either Alexa568-tubulin or mCherry-Lis1 was excited at 561 nm, alternating with simultaneous excitation of GFP-dynein at 488 nm and either Atto647N-tubulin, Alexa647-labelled SNAP-BicD2-N or Alexa647-labelled SNAP-EB3 at 647 nm.

### Analysis of dynein motility

To analyse DDB motion (Fig. EV3 and Fig. 5F-G), kymographs were generated from image sequences using the KymographClear 2.0 ImageJ plugin (Mangeol et al., 2016) (www.nat.vu.nl/∼erwinp/downloads.html). First, a maximum intensity projection image was generated that was used to define tracks using the segmented line tool. After drawing a single track, the plugin generated three distinct kymographs by Fourier filtering (Mangeol et al., 2016) showing separately forward moving, backward moving and pausing, or static particles. The trajectories in the kymographs were further analysed with the KymographDirect software (Mangeol et al., 2016). Spurious trajectories were rejected manually and fragmented trajectories from a single track were linked manually using ‘link’ option of the KymographDirect. Trajectories of directionally moving, diffusive or static fluorescent particles are identified by this software using the previously generated Fourier filtered kymographs. Statistical analysis of the velocities and run-lengths of the trajectories corresponding to directional motility was then performed automatically by the KymographDirect software.

To calculate the number of directionally motile events per μm of microtubule length (Fig. 5E), the total number of directionally motile events for each experimental condition was divided by the total length of dynamic microtubules along which the motile events occurred. The maximum lengths of dynamic microtubules were measured using the segmented line tool in ImageJ. The error bars represent the standard deviation.

### Quantification of fluorescence intensities at microtubule ends

To quantify and compare the fluorescence intensities of GFP-dynein at microtubule plus ends in end tracking experiments, averaged intensities profiles were generated as described previously (Roostalu et al., 2015). In brief, kymographs of dynamically growing microtubules were generated using ImageJ Multiple Kymograph plug-in for the GFP-dynein and fluorescently labelled microtubule channels. Then the microtubule plus ends were marked in the microtubule kymographs using the ‘segmented line’ tool in ImageJ. Kymographs of both channels were then computationally straightened using the marked microtubule plus end positions as a constant reference position. The fluorescence intensities at each position along all kymographs of a dataset were then averaged to generate time-averaged spatial intensity profiles for the two channels. For each assay condition the GFP-dynein intensity profiles show the average of mean intensities along a distance of 2 μm from three datasets. The error bars represent the standard deviation.

### Quantification of start probability of GFP-dynein run intiation on dynamic microtubule

To measure the probability of GFP-dynein run initiation on a dynamic microtubule, the distance of the start position of a run from the microtubule plus end was measured (initiation distance, Fig. 3A). The corresponding dynamic microtubule length at the time of initiation was also measured. Probabilities of initiation of DDB runs as a function of the distance from the microtubule plus end were calculated using Matlab. ‘1 - cumulative probability’ distribution functions (cdf) of initiation distances and of the corresponding microtubule lengths at the moment of initiation were computed (Fig. EV4 and Fig. EV6C). Histograms of relative spatial initiation probabilities within the first 7 μm from the microtubule plus end were extracted from the cumulative distributions, correcting for the measured microtubule length distribution (Fig. 3B, D, Fig. EV6B).

## ACKNOWLEDGEMENTS

We thank A. Carter for providing the plasmids for dynein Multibac generation, M. Steinmetz for EB3 plasmid, J. Gannon for cloning the EB3 expression construct, and N. Cade for help with image analysis. We thank the Cell Services Lab for the production of HeLa S3 cells, and the Proteomics Lab for mass-spectrometry of dynactin (both at the Francis Crick Institute). This work was supported by the Francis Crick Institute which receives its core funding from Cancer Research UK (FC001163), the UK Medical Research Council (FC001163), and the Wellcome Trust (FC001163). J.R acknowledges funding from a Sir Henry Wellcome Postdoctoral Fellowship (100145/Z/12/Z) and T.S. from the European Research Council (Advanced Grant, project 323042).

## AUTHOR CONTRIBUTIONS

RJ performed the experiments, analysed data. RJ, JR and TS designed the study and wrote the manuscript. MT cloned Lis1 and p150-N and transferred expertise.

## EXPANDED VIEW FIGURE LEGENDS

**Figure EV1 - related to Figure 1.**
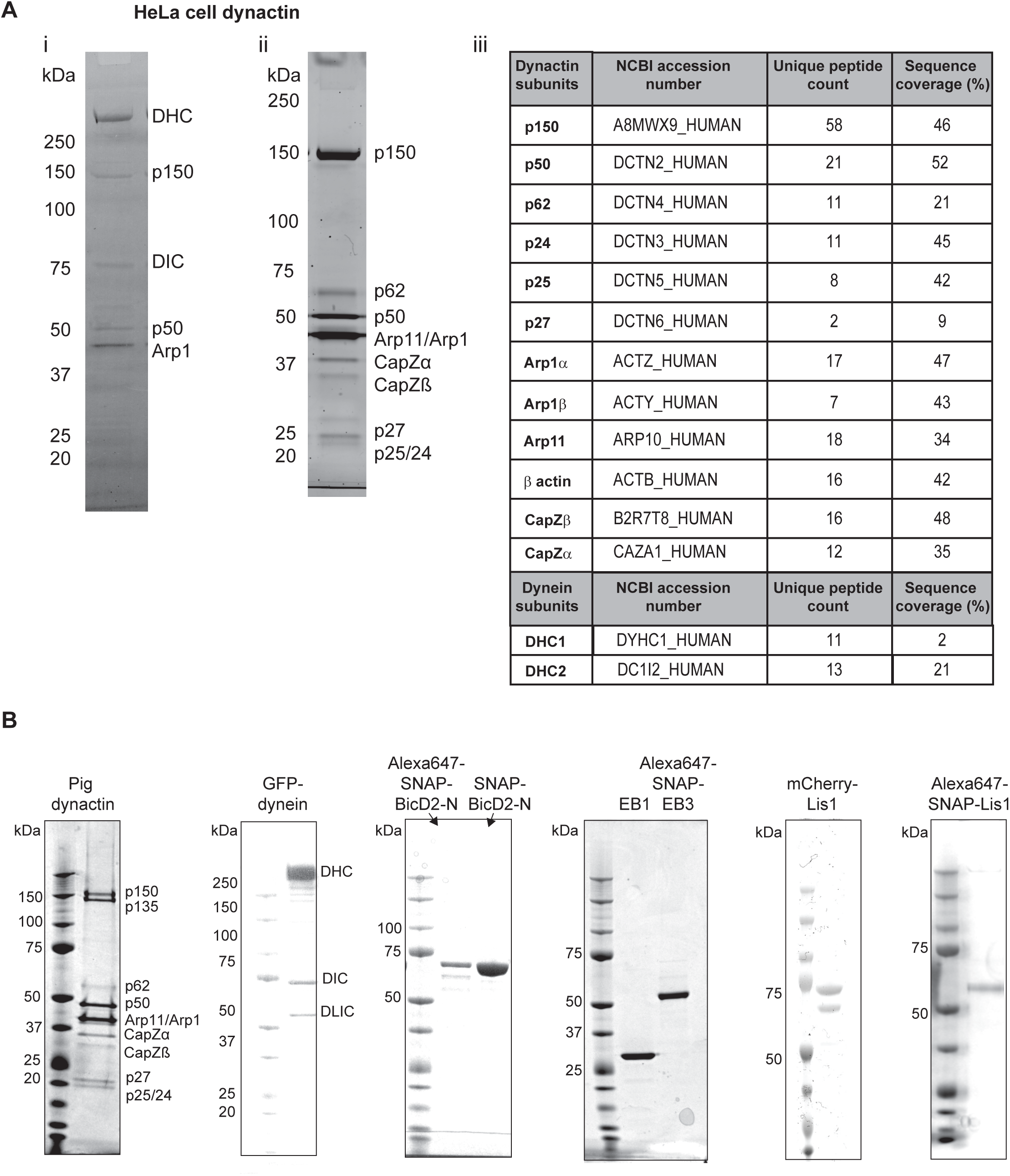
Purified proteins used in this study. (**A**) Purification of human dynactin from HeLa S3 cells: (i) Coomassie Blue-stained SDS-PAGE of the eluate from a BicD2 affinity column (first chromatography step of the dynactin purification). Bands corresponding to the dynein and dynactin complex are labelled. (ii) Sypro Ruby-stained SDS-PAGE of the pooled peak fractions of purified dynactin after anion exchange chromatography. (iii) Mass-spectrometry results of the purified human dynactin complex demonstrating the presence of all subunits of the dynactin complex; some fragments of the dynein heavy and intermediate chains, but not of the smaller dynein subunits were also detected. (**B**) Coomassie Blue-stained SDS PAGE of the purified proteins used in this study, as indicated. The individual dynactin and dynein subunits are labelled. (The double band of mCherry-Lis1 originates from different mCherry maturation states, as previously observed for other mCherry tagged proteins (Duellberg et al., 2014)).

**Figure EV2 - related to Figure 1.**
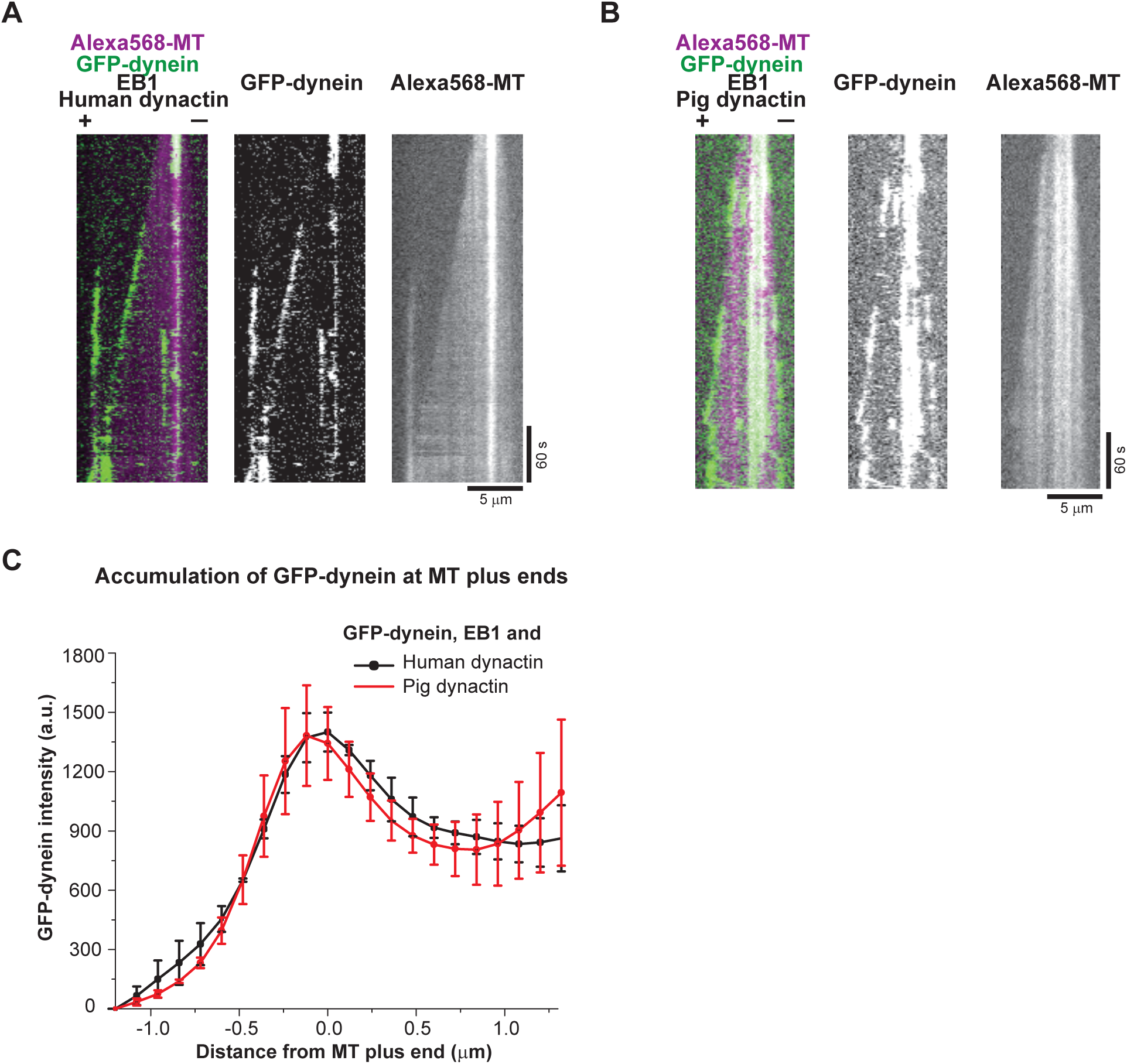
Microtubule end tracking of GFP-dynein in the presence of EB1, and either pig or human dynactin. (**A, B**) Dual colour kymographs of GFP-dynein (green) tracking the ends of Alexa568 microtubules (magenta in merge) in the presence of EB1 and either pig dynactin (A) or human dynactin (B). Protein concentrations were 10 nM GFP-dynein, 10 nM pig or human dynactin, 20 nM EB1, and 17.5 μΜ tubulin. (**C**) Averaged fluorescence intensity profiles of GFP-dynein at growing microtubule plus ends in the presence of EB1 and either human (black squares), or pig (red circles) dynactin. Mean values from three separate experiments (each with approximately 50 kymographs) are shown; error bars are s. d. Experiments were performed at 30°C.

**Figure EV3 - related to Figure 2.**
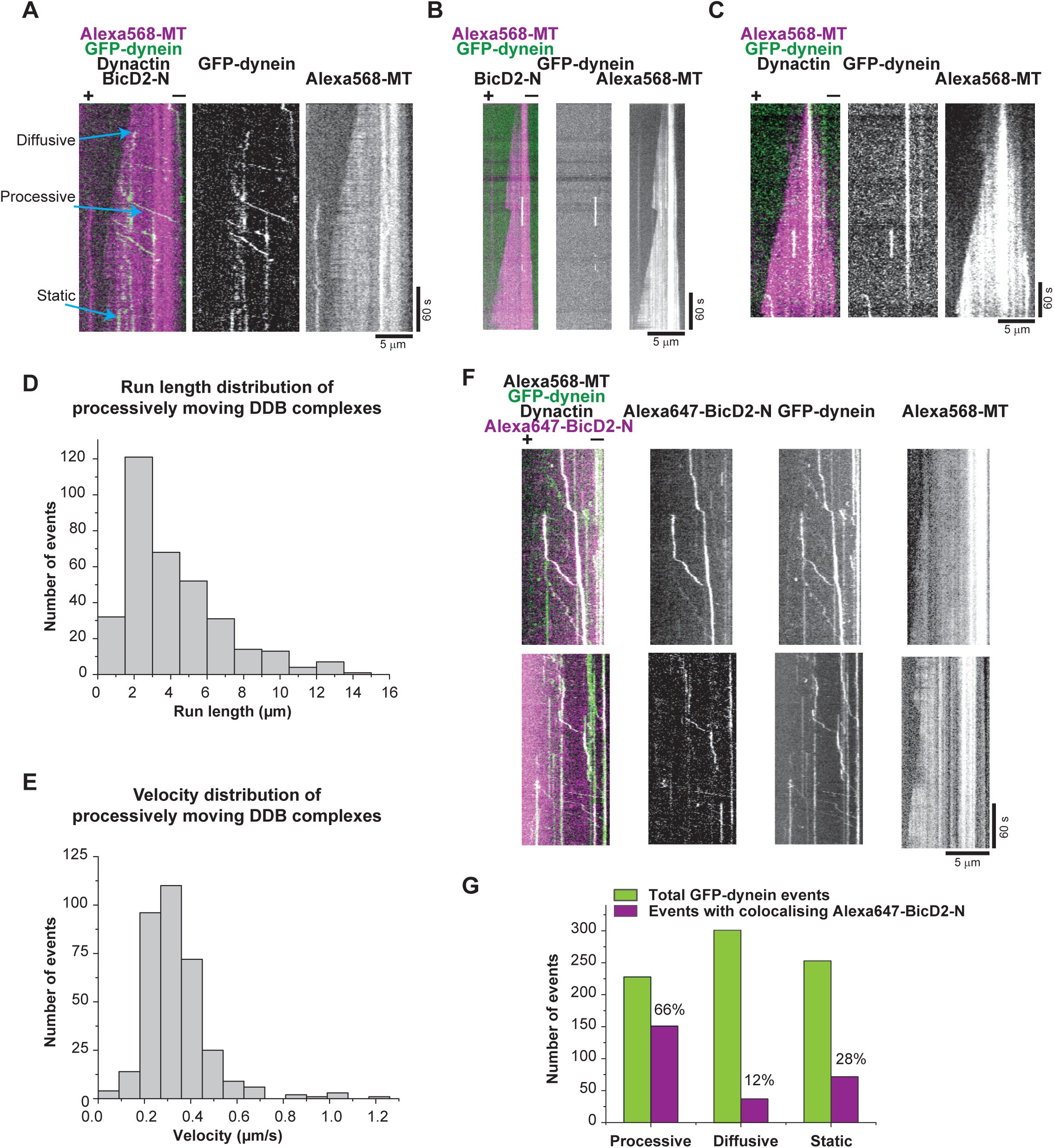
Dynein motility in the presence of dynactin and BicD2-N on dynamic microtubules. (**A**) Dual and single colour TIRF microscopy kymographs depicting GFP-dynein motility (green in merge) on dynamic Alexa568-microtubules (magenta in merge). Concentrations were 10 nM GFP-dynein, 20 nM dynactin, 200 nM BicD2-N, and 17.5 μΜ Alexa568-tubulin. (**B, C**) Kymographs showing GFP-dynein behaviour in the absence of **(B)** BicD2-N or **(C)** dynactin; other conditions as in A. (**D**) Histogram of the run length distribution of processively moving DDB complexes (n = 343 complexes from three separate experiments). The mean run length is 4.1 ± 0.15 (s.e.m) μm. (**E**) Histogram of the velocity distribution of processively moving DDB complexes (n = 343 complexes). The mean velocity is 0.34 ± 0.01 (s.e.m) μm/s. (**F**) Dual and single colour kymographs showing preferential colocalisation of GFP-dynein (green in merge) and Alexa647-BicD2-N (magenta in merge) during processive motility in the presence of dynactin on dynamic Alexa568-microtubules (not shown in merge). Protein concentrations were 10 nM GFP-dynein, 20 nM human dynactin, 200 nM Alexa647-BicD2, and 17.5 μM Alexa568-tubulin. Microtubule orientation as indicated. (**G**) Quantification of the number of observed GFP-dynein (green bars) and Alexa647-BicD2-N (magenta bars) events in the three categories ‘processive motility’, ‘diffusion’ and ‘static binding’ and the percentage of events with colocalising BicD2-N in each category. Most colocalisation was observed for processive motility events (n = 788 complexes from two separate experiments). Experiments were performed at 30°C.

**Figure EV4 - related to Figure 3.**
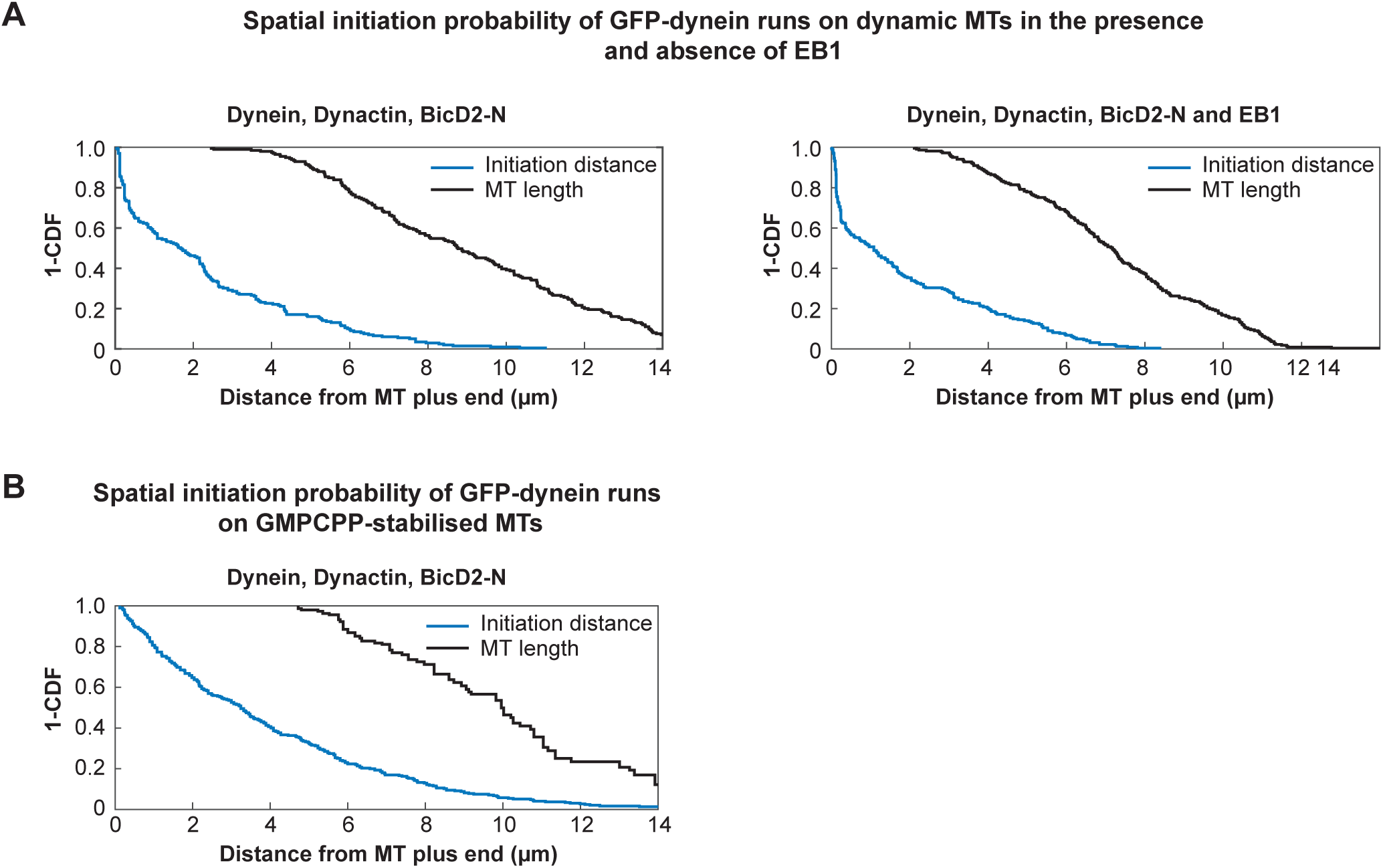
Distributions of initiation probabilities and microtubule lengths. ‘1 - cumulative probability’ distribution function of the distances of processive run initiation, measured from the microtubule plus end, and ‘1 - cumulative probability’ distribution function of the corresponding microtubule lengths measured at each moment of run initiation (black). Conditions as indicated (corresponding to conditions shown in Fig. 3B black, 3B red, and 3C).

**Figure EV5 - related to Figure 5.**
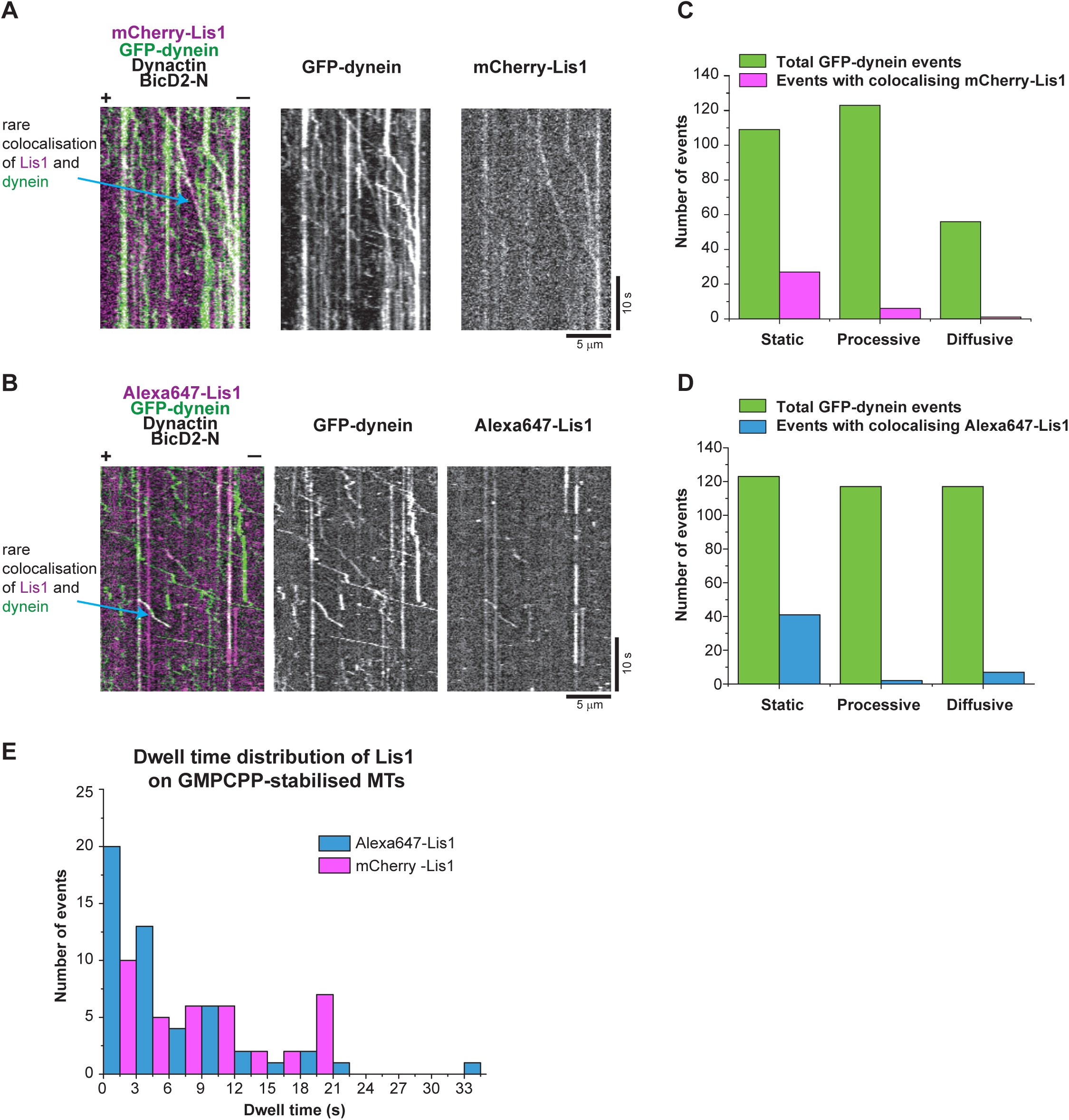
Colocalisation of GFP-dynein with Lis1. (**A**, **B**) TIRF microscopy kymographs depicting GFP-dynein (green) on GMPCPP stabilisied microtubules in the presence of dynactin, BicD2-N and either mCherry-Lis1 **(A)** or Alexa 647-Lis1 **(B)**. Rare colocalisation of dynein with Lis1 was observed. Microtubule plus and minus ends are labelled by (+) and (-). Concentrations were 10 nM GFP-dynein, 20 nM dynactin, 200 nM BicD2-N, 1 μΜ Lis1. (**C**, **D**) Quantification of the total number of observed GFP-dynein binding events (green bars) and those colocalising **(C)** with mCherry-Lis1 (magenta bars) or **(D)** with Alexa647-Lis1 (blue bars), separately counted for the three categories ‘processive motility’, ‘diffusion’ and ‘static binding’. The percentage of colocalisation events in each category is indicated. (**E**) Bar graph showing dwell time distribution of mCherryLis1 (magenta bars) and Alexa647-Lis1 (blue bars). Mean dwell time for Alexa647-Lis1, 8.61 ± 1.21 (s.e.m.) s, and for mCherry-Lis1, 8.59 ± 1.05 (s.e.m.) s. Experiments were performed at 30°C.

**Figure EV6 - related to Figure 5.**
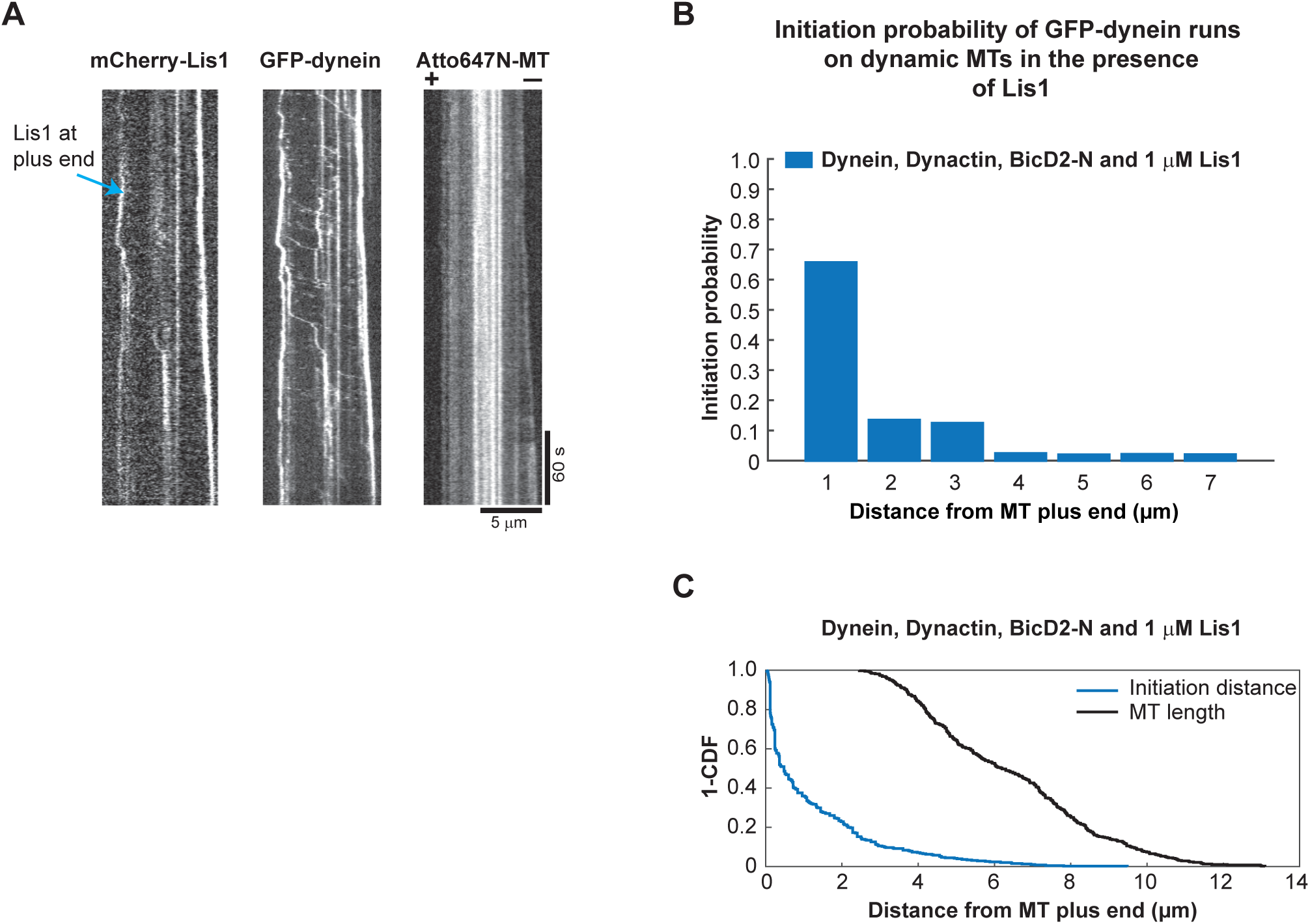
Spatial initiation probability of DDB runs within the first 7 μm from growing microtubule plus ends in the presence of EB1 and Lis1. (**A**) Single colour kymograph of a triple colour TIRF microscopy experiment showing mCherry-Lis1 co-localising with GFP-dynein at dynamic Atto647N-microtubule plus ends. Protein concentrations were as in Fig. 3B red with the additional presence of 1 μM mCherry-Lis1. Microtubule orientation as indicated. Experiments were performed at 30°C. **(B)** Histogram of spatial initiation probabilities of DDB runs in the presence of EB1 and Lis1. **(C)** Corresponding ‘1 - cumulabive probability’ distribution functions of microtubule lengths at each initiation moment and of the initiation probabilities. Over 200 complexes were analysed from three different data sets.

## MOVIE LEGENDS

**Movie 1.** Microtubule plus-end tracking of GFP-dynein (green) localising to the plus ends of dynamic Alexa568-microtubules (magenta) in the presence of dynactin and EB1. Experimental condition as in Fig. 1A.

**Movie 2.** GFP-dynein (green) on dynamic Atto565-microtubules (magenta) in the presence of all DDB components and EB1, showing DDB motion and reduced end-tracking of GFP-dynein. Experimental condition as in Fig. 2D.

**Movie 3.** Dynactin-mediated GFP-dynein (green) plus end tracking on Atto565-microtubules (magenta) at an elevated dynactin concentration. Experimental condition as in Fig. 4A.

**Movie 4.** p150-mediated GFP-dynein (green) plus end tracking on Atto565-microtubules (magenta). Experimental condition as in Fig. 4B.

**Movie 5.** GFP-dynein (green) on dynamic Atto647N-microtubules (magenta) in the presence of all DDB components, EB1, and 1 μΜ mCherry-Lis1, showing high number of processive DDB runs. Experimental condition as in Fig. 5B.

**Movie 6.** GFP-dynein (green) on dynamic Atto647N-microtubules (magenta) in the presence of all DDB components, EB1, and 5 μΜ mCherry-Lis1, showing GFP-dynein tracking plus-ends. Experimental condition as in Fig. 5C.

